# NRF2 activation induces NADH-reductive stress providing a metabolic vulnerability in lung cancer

**DOI:** 10.1101/2022.08.31.506025

**Authors:** Tommy Weiss-Sadan, Maolin Ge, Addriaan de Groot, Alexander Carlin, Magdy Gohar, Hannah Fischer, Lei Shi, Ting-Yu Wei, Charles H. Adelmann, Tristan Vornbäumen, Benedkit R. Dürr, Mariko Takahashi, Marianne Richter, Junbing Zhang, Tzu-Yi Yang, Vindhya Vijay, Makiko Hayashi, David E. Fischer, Aaron N. Hata, Thales Papaginanakopoulos, Raul Mostoslavsky, Nabeel Bardeesy, Liron Bar-Peled

## Abstract

Multiple cancers regulate oxidative stress by activating the transcription factor NRF2 through mutation of its negative regulator KEAP1. NRF2 has been studied extensively in KEAP1-mutant cancers, however the role of this pathway in cancers with wildtype KEAP1 remains poorly understood. To answer this question, we induced NRF2 via pharmacological inactivation of KEAP1 in a panel of 50+ non-small lung cancer cell lines. Unexpectedly, marked decreases in viability were observed in >13% of the cell lines—an effect that was completely rescued by NRF2 ablation. Genome-wide and targeted CRISPR screens revealed that NRF2 induces NADH-reductive stress, through the upregulation of the NAD^+^-consuming enzyme ALDH3A1. Leveraging these findings, we show that cells treated with KEAP1 inhibitors or those with endogenous KEAP1 mutations are selectively vulnerable to Complex I inhibition, which impairs NADH oxidation capacity and potentiates reductive stress. Thus, we identify reductive stress as a metabolic vulnerability in NRF2-activated lung cancers.

## Introduction

To support their rapid proliferation, tumors must adapt their metabolism to an ever-growing list of cell intrinsic and extrinsic pressures (Cantor and Sabatini, 2012; Kerk et al., 2021; Martinez-Reyes and Chandel, 2021; Vander Heiden and DeBerardinis, 2017). One such pressure is the maintenance of redox homeostasis as a pre-requisite for tumor proliferation (Chandel and Tuveson, 2014; Chio and Tuveson, 2017). Tumors have biochemically rewired core metabolic pathways to maintain elevated levels of biosynthetic molecules and consequently generate increased reactive oxygen species (ROS) compared to normal cells (Gorrini et al., 2013; Irani et al., 1997; Lee et al., 1999; Vander Heiden et al., 2009). High levels of ROS modify nucleic acids, proteins and lipids—and can provoke lethal cellular effects through multiple mechanisms, including creating DNA damage, impairing function of mitochondria and other organelles and disrupting the integrity of cell membranes (Chio and Tuveson, 2017; Sies et al., 2022).

While redox imbalance in cancer cells has been investigated extensively in the context of oxidative stress, the converse of oxidative stress, reductive stress and in particular its impact on malignant cells is poorly understood (Xiao and Loscalzo, 2020). Reductive stress is induced by excessive levels of antioxidants and high concentrations of reduced nucleotide cofactors required for antioxidant and detoxification reactions. An overly reductive cell state (Gores et al., 1989; Korge et al., 2015; Manford et al., 2021; Manford et al., 2020; Narasimhan and Rajasekaran, 2015) can affect vital cellular processes such as oxidative protein folding in the ER (Yang et al., 2007). Indeed, recent studies have demonstrated that reductive stress is just as harmful to cell proliferation as oxidative stress (Goodman et al., 2020; Ho et al., 2017; Perez-Torres et al., 2017; Zhang et al., 2012) with high levels of NADH leading to disruption of de novo lipid, amino acid and nucleotide biosynthesis as a result of decreased electron acceptors (Birsoy et al., 2015; Diehl et al., 2019; Garcia-Bermudez et al., 2018; Gui et al., 2016; Li et al., 2022; Sullivan et al., 2015).

To counter oxidative stress, tumors rely on NRF2, the central transcriptional regulator of the antioxidant response (DeNicola et al., 2011; Igarashi et al., 1994; Itoh et al., 1997; Pillai et al., 2022; Sporn and Liby, 2012). Under conditions of low oxidative stress, NRF2 binds to KEAP1, a tumor suppressor and ROS sensing protein, that leads NRF2 to its rapid proteasomal degradation (Baird et al., 2014; Suzuki and Yamamoto, 2015). Under high ROS-levels, key ROS-sensing cysteines in the backbone of KEAP1 are modified resulting in the dissociation and nuclear translocation of NRF2 (Kobayashi et al., 2004; Zhang et al., 2004), and the consequent induction of several hundred genes involved in antioxidant response (Malhotra et al., 2010; Sporn and Liby, 2012). NRF2 is activated in many cancers, including 30% of non-small cell lung cancers (NSCLCs) via genetic inactivation of KEAP1 (Cancer Genome Atlas Research, 2014). In *KEAP1*-mutant cancers, NRF2 is absolutely required for tumor growth functioning as a potent oncogene (Bar-Peled et al., 2017; DeNicola et al., 2011; Rojo de la Vega et al., 2018; Romero et al., 2017; Satoh et al., 2013).

To date, a significant body of our knowledge about the role of the NRF2-KEAP1 pathway in cancer comes from the discovery that NRF2 functions as an oncogene in the context of KEAP1-mutations in lung cancer (Chio et al., 2016; DeNicola et al., 2015; Fan et al., 2017; Goldstein et al., 2016; Leinonen et al., 2014; Romero et al., 2017; Sporn and Liby, 2012). We hypothesized that NRF2 activity might benefit the proliferation of NSCLCs when KEAP1 is not mutated. By activating NRF2 in a panel of NSCLCs we found, surprisingly, that rather than promote cell proliferation, activation of NRF2 via acute pharmacologic or genetic inhibition of KEAP1 potently blocks the growth of >13% of NSCLC cell lines –characterizing them as KEAP1-dependent. This dependency stems from a cell’s intrinsic preference for glucose utilization– with KEAP1-dependent cells characterized by lower levels of glycolysis and sensitivity to Complex I inhibitors. Mechanistically, we find that NRF2 activation results in an increase in NADH levels in KEAP1-dependent but not KEAP1-independent cells. We demonstrate through manipulation of glycolysis and NADH oxidation rates, that NADH levels are both necessary and sufficient to mediate sensitivity to NRF2 activation and uncover that ALDH3A1, a dehydrogenase involved in antioxidant response, has a primary role in increasing NADH levels and sensitizing KEAP1-dependent cells to NRF2 activation. Finally, we demonstrate that increased NADH levels due to treatment with KEAP1-inhibitors or the presence of KEAP1-mutations confers exquisite sensitivity to a clinical grade Complex I inhibitor, which increases NADH levels and overwhelms NADH homeostasis in these cells, leading to reductive stress. Thus, we reveal how over-activation of an antioxidant signaling pathway leads to a reduced cellular state that can create “oxidative addiction” and synthetic lethal opportunities within a subset of lung cancers.

## Results

### Identification of KEAP1-dependent NSCLC cell lines

Multiple studies have identified a proliferation benefit from NRF2 activation in the context of KEAP1-mutant NSCLC cell lines (Bar-Peled et al., 2017; DeNicola et al., 2015; Romero et al., 2017; Singh et al., 2006). We hypothesized that additional NSCLC cell lines that are KEAP1-wildtype (WT) would also gain a proliferative advantage following NRF2 activation. To test this hypothesis, we treated a panel of 50+ genetically diverse NSCLC cell lines (with a majority of lines WT for KEAP1, NRF2 and CUL3) with KI696, a potent and specific inhibitor of KEAP1-NRF2 interactions (Davies et al., 2016; Ding et al., 2021), which leads to NRF2 stabilization and activation (Figure 1A). Proliferation was not altered in most lines, including *KEAP1*-mutants, (e.g., H2122) and was increased in a limited subset (Figure 1B). Surprisingly, we find that >13% of NSCLC cell lines have a substantial block in proliferation following KEAP1-inhibition (Figure 1B). In support of NRF2 activation mediating the proliferation block with KI696, we found that treatment of a subset of NSCLC lines with bardoxolone, a well-established NRF2 activator, also potently blocked proliferation (Figures S1A-B). Expression of KEAP1-targeting sgRNAs or doxycycline (DOX)-inducible shRNAs strongly blocked the proliferation of KI696-sensitive cell lines in comparison to non-targeting sgRNA/shRNAs, characterizing these cells as KEAP1-dependent (Figures 1C, S1C-E). Proliferation was not impacted in KI696-insensitive cells following genetic depletion of KEAP1, marking them as KEAP1-independent (Figures 1C, S1C-E). KEAP1-dependence could be rescued by re-expressing a sgKEAP1-mutant KEAP1 cDNA under a DOX-repressible element, and we observed a proliferation block in KI696-sensitive lines following DOX treatment and subsequent depletion of KEAP1 (Figures S1F-G).

**Figure 1:**
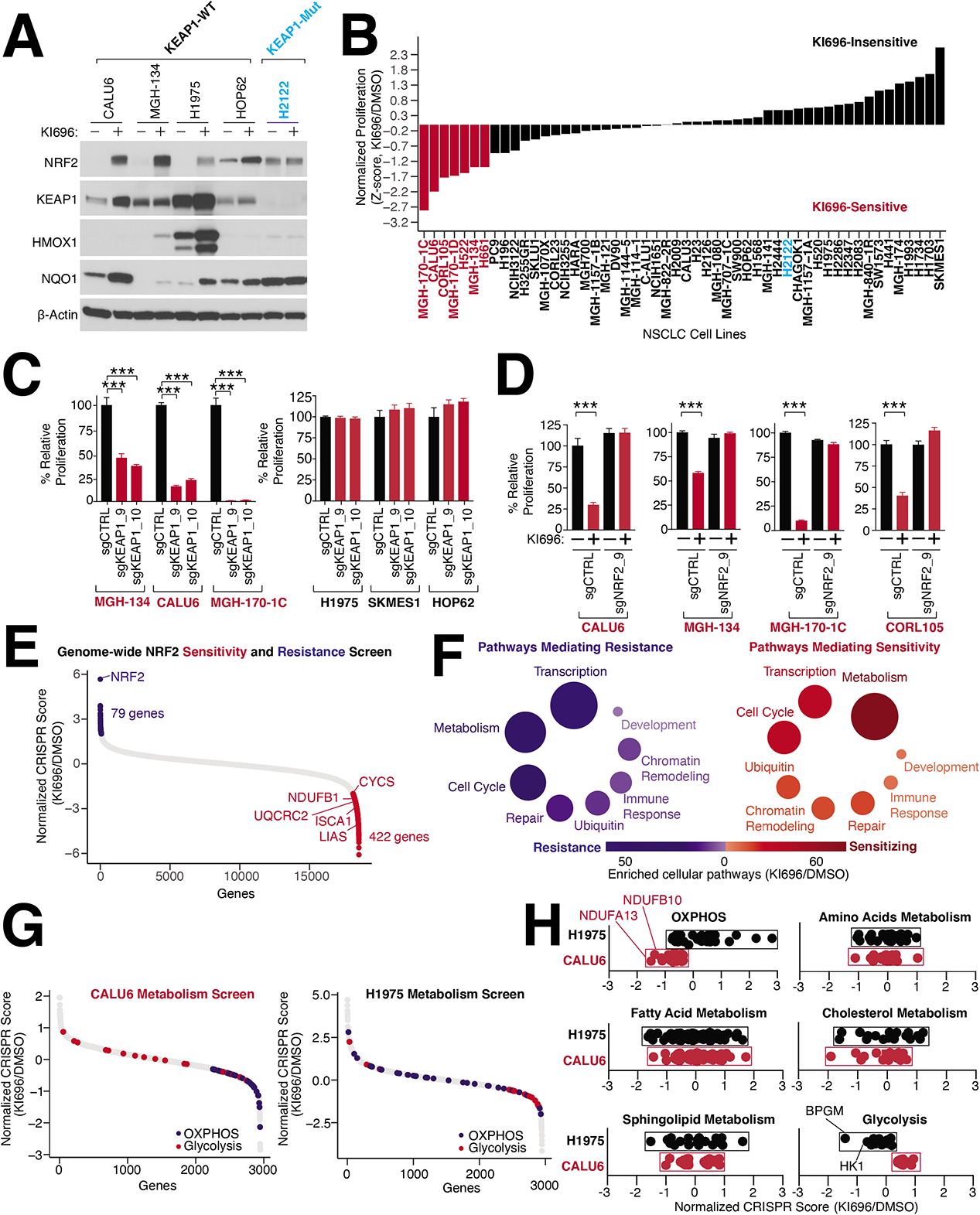
Identification of KEAP1-dependent NSCLC cell lines. (A) Representative immunoblot analysis of NSCLC cell lines following treatment with the NRF2-activator KI696 (1μM) for 48 hrs. (B) NRF2 activation in a panel of 50+ NSCLC cell-lines identifies KI696-sensitive cell lines. Cells were pre-treated with KI696 (1 µM) for 48 hrs and proliferation was determined by crystal violet staining following another 6 days of treatment. (C) Depletion of KEAP1 blocks proliferation of KI696-sensitive cell lines. KI696-sensitive (herein referred to as KEAP-dependent) and KI696-insensitive (KEAP1-independent) cell lines expressing sgRNAs targeting KEAP1 or a non-targeting control were analyzed for proliferation defects as described in (B) (Data are represented as a mean ± SEM, n=5 biological replicates). (D) Depletion of NRF2 rescues KI696-sensitivity. NSCLC cell lines expressing the indicated sgRNAs were treated with KI696 (1 µM) and proliferation was determined as described in (B) (Data are represented as a mean ± SEM, n=5 biological replicates). (E) Genome-wide CRISPR screen identifies genes mediating resistance and sensitivity to KEAP1-dependency. Highlighted genes are key mediators of sensitivity (red) or resistance (blue) (see also **Supplementary Table 1**). (F) Genes localized to metabolic pathways function as key mediators of sensitivity to NRF2 activation in CALU6 cells (see also **Supplementary Tables 2-3**). (G) Metabolism-focused CRISPR screen identifies metabolic regulators of NRF2 sensitivity. KEAP1-dependent CALU6 (red) and KEAP1-Independent (black) H1975 cells were infected with a metabolism focused sgRNA library and treated as described in (E) (see **also Supplementary Tables 4 and 5**). (H) Inactivation of oxidative phosphorylation (OXPHOS) sensitizes KEAP1-dependent cells to NRF2 activation whereas blockage of glycolysis sensitizes KEAP1-independent cells to NRF2 (see also Figure S2A). Statistical significance was determined by One-way ANOVA with Sidak’s corrections for multiple comparisons. *** indicates *p*-values < 0.0001.

The strong reliance on KEAP1 for proliferation was quite unexpected given the canonical characterization of KEAP1 as a recurrently mutated tumor suppressor in lung cancer (Cancer Genome Atlas Research, 2014; Kerins and Ooi, 2018; Leinonen et al., 2014; Romero et al., 2017; Sanchez-Vega et al., 2018; Singh et al., 2006; Sporn and Liby, 2012; Wu and Papagiannakopoulos, 2020). In KEAP1-dependent cells, KI696 treatment leads to a stabilization of NRF2 and expression of NRF2 target genes (Figures 1A, S1A) suggesting that KEAP1-dependency is associated with NRF2 activation. Depleting NRF2 in four KEAP1-dependent cell lines did not alter proliferation at baseline (Figures 1D, S1H), but completely reversed the proliferation arrest following pharmacological or genetic inhibition of KEAP1 (Figures 1D, S1H-J). By analyzing cancer essentiality data from genome-wide CRISPR screens across 800+ cancer cell lines (Lenoir et al., 2018; Tsherniak et al., 2017) we find that multiple cancer cell lines of different origins are sensitive to loss of KEAP1, including breast and skin cancers which share a similar rate of dependency as NSCLCs (Figure 1K). These results illustrate that KEAP1-dependency is broadly observed in multiple cancer subtypes highlighting the complex role of established oncogenes when their corresponding tumor suppressors are not mutated (Davoli et al., 2013; Serrano et al., 1997).

### Functional genomic interrogation of KEAP1 dependency

We did not find a correlation between KEAP1-dependency and mutational status of other oncogenic pathways (e.g., LKB1, PI3K, p53, KRAS). To search for genes which may predict sensitivity or resistance to NRF2 activation, we performed a genome-wide CRISPR screen in the KEAP1-dependent cell line CALU6. Following infection with the sgRNA library (10 sgRNAs/gene), cells were grown for 11 population doublings in the presence of 1 µM of KI696 or vehicle control. For each gene, we calculated a CRISPR score by comparing the relative fold change between corresponding sgRNAs enriched in KI696 vs. vehicle. This analysis identified 79 genes mediating resistance and 422 genes mediating sensitivity to KI696 (Figure 1E). Validating our screening approach, we identified NRF2 as the top scoring gene that mediated resistance when depleted in KEAP1-dependent cell lines (Figure 1E). There was also a striking enrichment of metabolic genes identified as mediators of KI696 sensitivity in the CRISPR screen, including many genes belonging to mitochondrial metabolic pathways (Figure 1F, **Supplementary Tables 1-3**). To better define metabolic mechanisms of sensitivity and resistance to NRF2 activation, we next undertook a metabolism-focused CRISPR screen, encompassing ∼2000 genes connected to different metabolic processes (Birsoy et al., 2015) in KEAP1-dependent (CALU6) and KEAP1-independent (H1975) cells following KI696 treatment (Figure 1G**, Supplementary Tables 4-5**). We found an enrichment for a subset of glycolytic genes mediating sensitivity in KEAP1-independent cells, whereas those genes enriched in oxidative phosphorylation (OXPHOS) sensitized KEAP1-dependent cells (Figures 1H, S2A). The absence of additional glycolytic or OXPHOS genes scoring in this screen most likely stems from their pan-essential nature.

### Anaerobic metabolism mediates resistance to NRF2 activation

To further investigate the metabolic basis for KEAP1-dependency, we focused on the opposing forms of glucose utilization that were differentially essential following KI696 treatment in KEAP1-dependent and –independent cells. The ratio of extracellular acidification rate (ECAR) to oxygen consumption rate (OCR) is used to characterize the preference for aerobic vs anaerobic metabolism (Dar et al., 2017). By comparing the OCR/ECAR ratio of 12 NSCLC models to their corresponding sensitivity for KI696, we found a strong correlation between a preference for aerobic metabolism and NRF2 sensitivity, whereas cells with high glycolytic rates were insensitive to NRF2 activity (Figure 2A). These results were further substantiated by comparing KEAP1-dependency to a glycolytic gene signature, with NSCLCs marked by a high glycolytic gene signature having a lower sensitivity to NRF2 activation (Figure 2B**, Supplementary Table 6**). Interestingly, we found that KEAP1-independent cells had on average higher lactate dehydrogenase (LDH) activity compared to their dependent counterparts (Figure S2B). Moreover, KEAP1-independent cell lines had higher levels of phosphofructokinase (PFK) one of the rate-limiting enzymes in glycolysis (Figures 2C, S2C). The higher glycolytic rates characterizing KEAP1-independent cells suggested this form of glucose utilization might be a mechanism to adapt to NRF2 activation. Therefore, we over-expressed PFK in KEAP1-dependent CALU6 cells, finding not only an increase in glycolysis as measured by lactate secretion but a partial rescue of proliferation following NRF2 activation (Figures 2D, S2D-E).

**Figure 2:**
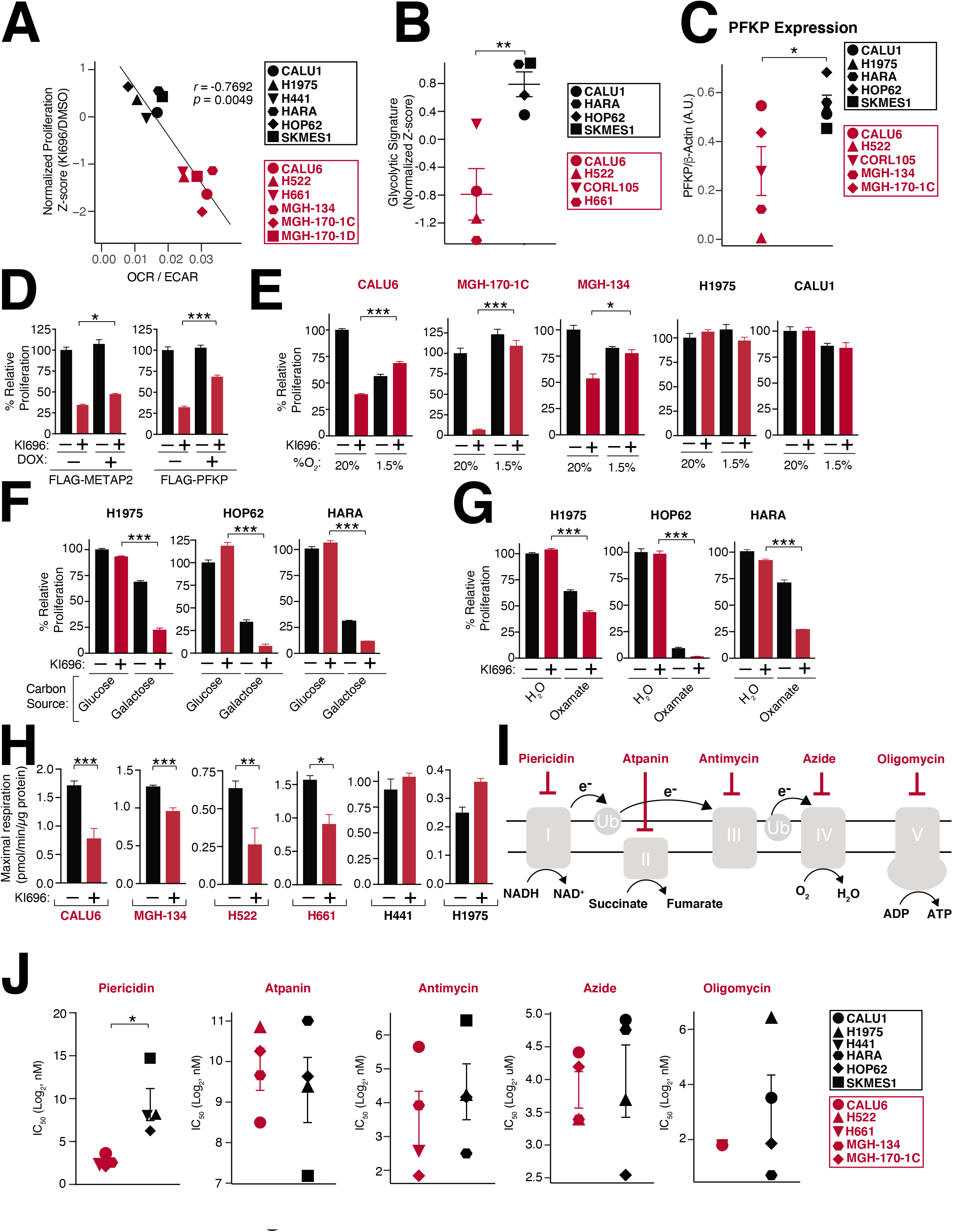
Metabolic requirements for NRF2 sensitivity. (A) NRF2 sensitivity correlates with higher levels of oxidative metabolism. Oxygen consumption rate (OCR) and extracellular acidification (ECAR) were measured in a panel of NSCLC cells and the OCR/ECAR for each cell line was plotted against its corresponding sensitivity to NRF2 activation (Data are represented as a mean ± SEM, n=6-8 biological replicates). (B) KEAP1-dependent cells have a lower glycolytic gene signature (see also **Supplementary Table 6**). (C) The rate-limiting glycolytic enzyme phosphofructokinase, platelet isoform (PFKP), is highly expressed in KEAP1-dependent cells. Quantification of PFKP levels relative to β-actin (see also Figure S2B). (D) PFKP overexpression restores proliferation following NRF2 activation. Relative proliferation of CALU6 cells expressing FLAG-PFKP or FLAG-METAP2 (control) was determined by crystal violet staining following doxycycline (DOX) (100 nM) and KI696 (1 µM) treatment (Data are represented as a mean ± SEM, n=5 biological replicates). (E) Hypoxia rescues NRF2 sensitivity. Relative proliferation in a panel of NSCLC cell lines following treatment with KI696 (1 µM) and culture in normoxic (20% O_2_) or hypoxic conditions (1.5% O_2_) was determined as in (D) (Data are represented as a mean ± SEM, n= 5 biological replicates). (F-G) Glycolytic inhibition sensitizes cells to NRF2 activation. KEAP1-independent cells were treated with KI696 (1 µM) and cultured in media containing glucose (10 mM) or galactose (10 mM) (F) or co-treated with sodium oxamate (10 mM) (G) and relative proliferation was determined as in (D) (Data are represented as a mean ± SEM, n=5 biological replicates). (H) NRF2 activation decreases maximal respiration in KEAP1-dependent cells. Maximal respiration was determined in a panel of NSCLC lines following treatment with KI696 (1 µM) for 48 hrs (Data are represented as a mean ± SEM, n=6-8 biological replicates) (see also Figure S3A). (I-J) Complex I inhibition is selectively toxic to KEAP1-dependent cell lines. Schematic of different ETC inhibitors used in this study (I). IC_50_-values (J) were determined for a panel of NSCLC cell lines (Data are represented as a mean ± SEM, n= 4-5 samples/group measured in 4-6 biological replicates) (see also Figure S3C). * indicates *p*-values < 0.05, *** indicates *p*-values < 0.0001. One-way ANOVA with Sidak’s post-hoc correction and two-tailed student’s t-test were used to determine statistical significance.

Given the well-established connection between hypoxia and glycolytic reprogramming (Tennant et al., 2009), we induced hypoxia in KEAP1-dependent cells, finding an increased expression in the glycolytic enzyme HK2 (Figure S2F). Importantly, induction of hypoxia largely rescued NRF2-sensitivity in KEAP1-dependent cells while having no effect on KEAP1-independent cells (Figure 2E). Our results suggest that high glycolytic rates are sufficient to overcome NRF2 sensitivity and we asked whether decreasing glycolysis within KEAP1-independent cells results in NRF2-sensitivity. To this end, we blocked the initial and terminal steps of glycolysis by growing KEAP1-independent cells in galactose-containing media or treating cells with sodium oxamate a LDH inhibitor (Zhai et al., 2013), respectively. As expected, both treatments resulted in a strong sensitization to NRF2 activation in KEAP1-independent cells (Figures 2G-H). These results indicate that upregulation of glycolysis is both necessary and sufficient to rescue NRF2 sensitivity.

### NRF2 activation disrupts mitochondrial metabolism in KEAP1-dependent cells which are hypersensitive to Complex I inhibition

We next investigated the impact of NRF2 activation on mitochondrial respiration (Sayin et al., 2017), finding a substantial decrease in maximal respiratory capacity in KEAP1-dependent cells following KI696 treatment (Figures 2H, S3A). This defect in respiration was the result of NRF2, as loss of the transcription factor completely rescued OCR following KI696 treatment (Figure S3B). Inhibition of mitochondria function also extended to mitochondrial metabolism, where we found that TCA metabolites were largely downregulated following NRF2 activation in KEAP1-dependent cell lines and is consistent with previous studies (Ding et al., 2021; Sayin et al., 2017) (Figure S3C, **Supplementary Table 7**). Importantly, KI696-mediated defects in TCA metabolism are completely dependent on NRF2 in CALU6 cells (Figure S3D, **Supplementary Table 8**). Curiously, we did not observe substantial changes in mitochondrial gene or protein expression in KEAP1-dependent cells following NRF2 activation (Figures S4A-B, **Supplementary Tables 9-10**) in comparison to canonical NRF2 targets, suggesting NRF2-regulation of mitochondrial function is post-translational or dependent on a change in metabolite(s) levels. These results indicate that NRF2 activation can be particularly detrimental to cancer cells which are reliant on mitochondrial function for their proliferation.

Given the prominent role of the electron transport chain (ETC) in regulating oxygen consumption, we asked if there was altered sensitivity to inhibition of ETC complexes between KEAP1-dependent and KEAP1-independent cells. In general, we found comparable IC_50_ values for ETC inhibitors targeting complexes 2-5 between dependent and independent cells. However, KEAP1-dependent cells demonstrated a striking sensitivity to the Complex I inhibitor piericidin (Figures 2I-J). To further confirm the reliance of KEAP1-dependent cells on Complex I, we tested additional inhibitors, including phenformin and rotenone, again finding on average lower IC_50_ values for KEAP1-dependent cells in comparison to their independent counterparts (Figure S4D). Collectively, these results establish that a cell’s intrinsic preference for glucose utilization (aerobic vs aneorobic fermentation) dictates its sensitivity to NRF2 activation, with inhibition of Complex I, a particular liability for KEAP1-dependent cells.

### NRF2 activation results in NADH reductive stress in KEAP1-dependent cells

We hypothesized that NRF2 sensitivity arises from a shared metabolic activity in KEAP1-dependent and –independent cells and only reveals itself upon NRF2 activation because it is limiting in dependent cells but not in independent cells. Our metabolic characterization of KEAP1-dependence revealed a heightened sensitivity to Complex I inhibition and that an increase in glycolytic rates could reverse NRF2 sensitivity. Because Complex I activity and glycolysis have the shared activity of NADH oxidation, we investigated whether NRF2 activation results in heightened NADH levels, which can block multiple cellular reactions dependent on NAD^+^ (Birsoy et al., 2015; Diehl et al., 2019; Garcia-Bermudez et al., 2018; Gui et al., 2016; Li et al., 2022; Sullivan et al., 2015). We constructed a panel of NSCLC cell lines that stably expressing SoNar, a genetically encoded NADH/NAD^+^ reporter (Zhao et al., 2015) that measures differences in the relative levels of NADH and NAD^+^. At baseline, we found that KEAP1-dependent and KEAP1-independent cells had a similar ratio of NADH/NAD^+^ (Figure S5A). However, following KI696 treatment we found a consistently higher NADH/NAD^+^ ratio generated in KEAP1-dependent cells compared to KEAP1-independent cells that was mirrored by traditional methods to measure NADH/NAD^+^ (Figure 3A, S5B-D). Furthermore, genetic depletion of KEAP1 using DOX-inducible shRNAs also mediated an increase NADH/NAD^+^ ratio in KEAP1-dependent but not KEAP1-indendent cells (Figures S5E-I). The increase in NADH/NAD^+^ ratio was completely reliant on NRF2, as depletion of the transcription factor blocked the KI696-mediated increase in the NADH/NAD^+^ ratio (Figures 3B, S5J). These results demonstrate NRF2 activation is both necessary and sufficient to mediate high NADH levels in KEAP1-dependent cells.

**Figure 3:**
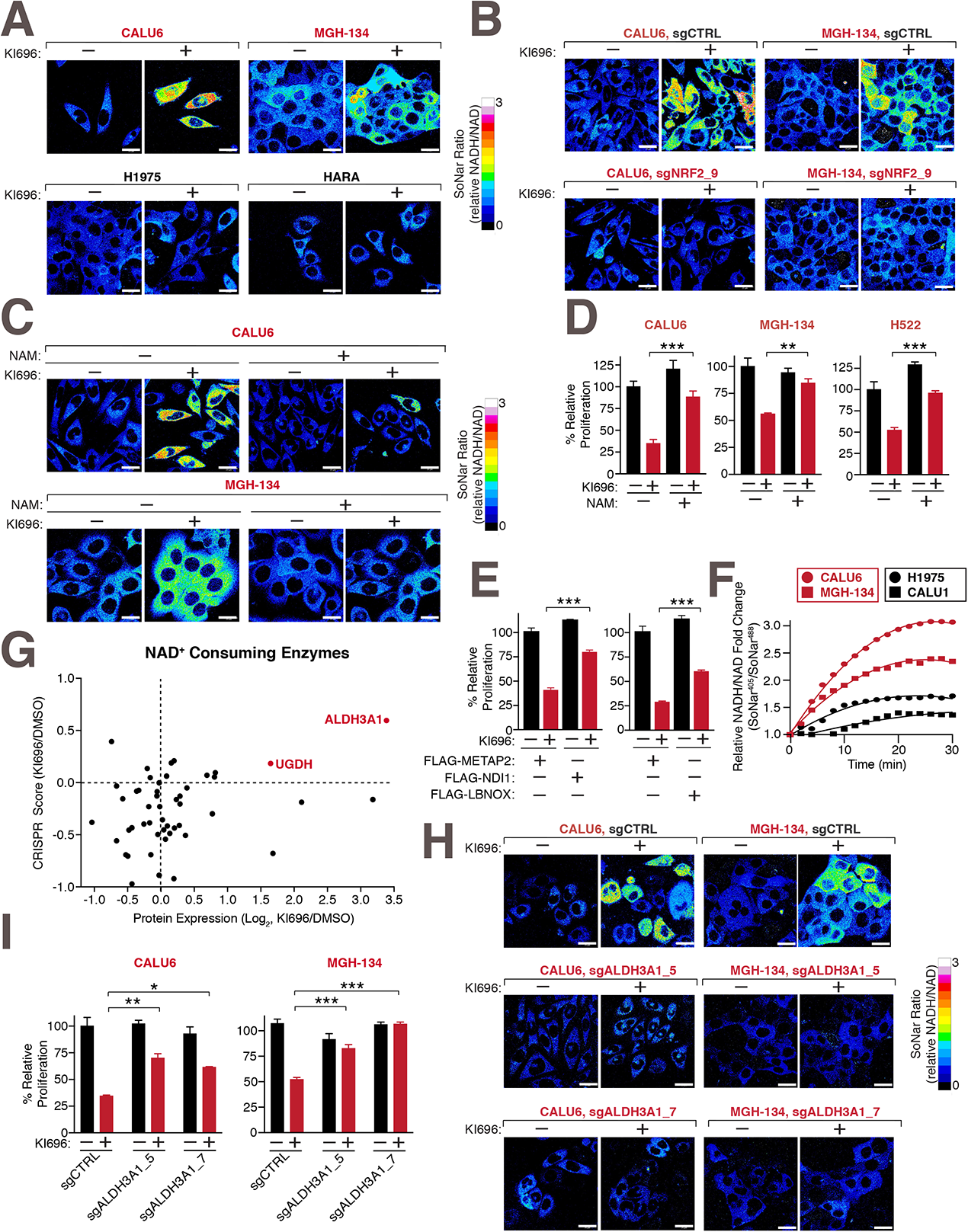
NRF2 induces NADH-reductive stress in KEAP1-dependent cells. (A) NRF2 activation increases the NADH/NAD^+^ ratio in KEAP1-dependent but not KEAP1-idependent cells. Immunofluorescence analysis of NSCLC cell lines stably expressing the NADH/NAD^+^ reporter SoNar following treatment with KI696 (1 µM) for 48 hrs. The representative NADH/NAD^+^ ratiometric image was constructed by taking the ratio of the emission intensity of 408 (NADH binding) vs 488 (NAD^+^ binding) for SONAR (see also Figures S5C-D). (B) NRF2 depletion rescues KI696-mediated NADH/NAD^+^ increase. NSCLC cells expressing SoNar as in (A) and corresponding sgRNAs targeting NRF2 or a control were treated with KI696 and cells were analyzed as in (A) (see also Figure S5J). (C-D) Supplementation with NAM restores the NADH/NAD^+^ ratio following NRF2 activation and rescues proliferation in KEAP1-dependent cells. KEAP1-dependent NSCLCs were treated with KI696 and NAM (1mM) where indicated and analyzed as in (A) or assayed for a change in proliferation by crystal violet staining 6 days post treatment (D) (Data are represented as a mean ± SEM, n= 4-5 biological replicates) (see also Figure S6A). (E) Over-expression of NADH oxidizing enzymes partially rescues NRF2 activation. CALU6 cells stably expressing NDI1, LbNOX or METAP2 (control) were treated with KI696 and assayed for proliferation as described in (D) (Data are represented as a mean ± SEM, n= 5 biological replicates). (F) KEAP1-dependent cells have a higher rate of Complex I NADH oxidation compared KEAP1-independent NSCLCs. NSCLC cell lines stably expressing SoNar were treated with rotenone (0.5 µM) and analyzed by flow cytometry taking the ratio of the emission intensity at λem 530nm after excitation at λex 408nm (NADH binding) or λex 488nm (NAD^+^ binding). (G) Expression and dependency of NAD^+^ consuming enzymes following NRF2 activation in CALU6 cells. Scatter plot of NAD^+^-consuming enzymes identified in metabolism focused CRISPR screen (see also **Supplementary Table 4**) and proteomics (see also **Supplementary Table 10**) following KI696 treatment. (H) ALDH3A1 depletion rescues high NADH levels following NRF2 activation. CALU6 and MGH-134 cells stable expressing SoNar and the indicated sgRNAs were treated with KI696 and analyzed as described in (A) (see also Figure S6F). (I) Loss of ALDH3A1 rescues proliferation following NRF2 activation. KEAP1-dependent NSCLC cells expressing the indicated sgRNAs were treated with KI696 (1 µM) and proliferation was determined as described in (D) (Data are represented as a mean ± SEM, n= 5 biological replicates per condition). * indicates *p*-values < 0.05, ** indicates *p*-values < 0.01, *** indicates *p*-values < 0.0001. One-way ANOVA with Sidak’s post-hoc correction used to determine statistical significance. Scale bar: 25 µm.

To determine whether high levels of NADH were a primary mechanism underlying NRF2-sensitivity, we treated cells with β-nicotinamide mononucleotide (NAM), a NAD precursor (Micheli et al., 1990), which substantially rescued both high NADH/NAD^+^ ratio upon NRF2 activation and the concomitant proliferation block in KEAP1-dependent cells (Figures 3C-D, S6A). Supplementation with NAM reversed the KI696-mediated defects in respiration in KEAP1-dependent cells (Figure S6B), most likely by providing the required NAD^+^ equivalents for the TCA cycle. Furthermore, KEAP1-dependent cells expressing the NADH oxidizing enzymes LbNOX or NDI1 (McElroy et al., 2020; Seo et al., 1998; Titov et al., 2016) were partially protected from NRF2-mediated defects in proliferation and respiration in comparison to cells expressing a control protein (METAP2) (Figures 3E, S6C-D). Conversely, KEAP1-independent cells treated with a pyruvate kinase inhibitor leading to the activation of pyruvate dehydrogenase which increases the NADH/NAD^+^ ratio by shunting pyruvate away from lactate production (Luengo et al., 2021), resulting in an increased sensitivity to NRF2 activation (Figure S6E). These results suggested that upon activation of NRF2 in KEAP1-dependent cells, NADH-reductive stressed is induced leading to a proliferation block.

Our findings suggested that KEAP1-dependency may arise, in part, from a metabolic preference to utilize Complex I for NADH oxidation, resulting in a corresponding sensitivity to Complex I blockade previously observed. To provide evidence for this hypothesis, we measured NADH/NAD^+^ ratio in KEAP1-dependent and KEAP1-independent cells following sequential treatments with rotenone and oxamate. We found that KEAP1-dependent cells have a higher NADH/NAD^+^ ratio following Complex I inhibition compared to their independent counterparts (Figure 3F). However, LDH-based NADH oxidation, accounted for the majority of the NADH oxidation in both cell types and was indeed higher in KEAP1-independent cells (Figure S6F). Collectively, our findings provide one explanation by which cells with lower glycolytic rates are unable to cope with NADH reductive stress brought upon NRF2 activation.

### ALDH3A1 partially underlies NAHD-reductive stress in KEAP1-dependent cells

To explore the mechanisms by which NRF2 activation increases NADH levels in KEAP1-dependent cells, we focused on 105 enzymes that consume NAD^+^ and are up-regulated following NRF2 activation. We further filtered corresponding NAD^+^-consuming genes based on their ability to mediate resistance to NRF2 activation when depleted, identifying ALDH3A1 as a compelling candidate (Figure 3G). ALDH3A1 is a NRF2-responsive gene (Figures S4A-B) that functions in antioxidant defense, specifically the resolution of lipid peroxide stress by converting aldehydes to their corresponding carboxylic acids (Pappa et al., 2003; Wang et al., 2017). The dehydrogenase activity of ADLH3A1 results in the conversion of NAD^+^ to NADH and we found that depletion of this gene led to a substantial decrease in NADH levels following NRF2 activation in KEAP1-dependent cells (Figures 3H, S6G). Importantly, depletion of ALDH3A1 in KEAP1-dependent cells overcame the proliferation block following NRF2 activation (Figures 3I, S6H). These results illustrate how in the absence of oxidative stress, over-expression of antioxidant defense genes reshapes the cellular redox state to an overly reduced environment leading to a block in proliferation.

### Disruption of Complex I activity is a metabolic liability in KEAP1-mutant cells

Our results suggest that NRF2 can result in toxic levels of NADH, constituting a liability that may be exploited in cellular states with hyperactive NRF2 signaling. We tested this hypothesis by treating a panel of KEAP1-dependent, KEAP1-independent and KEAP1-mutant cells with IACS-010759 (IACS), a potent Complex I inhibitor (Molina et al., 2018), finding that KEAP1-mutant cells had a substantial increase in NADH levels following treatment (Figures 4A, S7A). Consistent with our hypothesis, KEAP1-independent cells, that were co-treated with KI696 and IACS also had substantial increases in NADH levels (Figures 4A, S7A). The increase in NADH levels following IACS treatment, directly translated to lower IC_50_ values in KEAP1-mutant and KEAP1-dependent cell lines compared to KEAP-independent cells (Figure 4B). Moreover, we found IACS potently blocked the growth of KEAP1-dependent and NRF2 activated cells in soft agar but not KEAP1-independent cells (Figures 4C, S7B-C). By depleting ALDH3A1 we found a partial rescue of IACS treatment in KEAP1-mutant cells (Figures 4D, S7D). Over-expression of LBNOX and NDI1 in KEAP1-mutant cells or treatment with NAM, also led to a partial rescue of IACS treatment (Figures 4E-F, S7E). These results reveal that NADH levels are at their tipping point in KEAP1-mutant cells and that a slight disruption in NADH oxidation through Complex I inhibition coordinated with hyper-active NRF2 signaling overwhelms NADH homeostasis leading to reductive stress and a blockage in cell proliferation (Figure 4G). Collectively, our results illustrate how reductive stress can be leveraged as a synthetic lethal opportunity within a genetically– and metabolically-defined subset of cancers.

**Figure 4:**
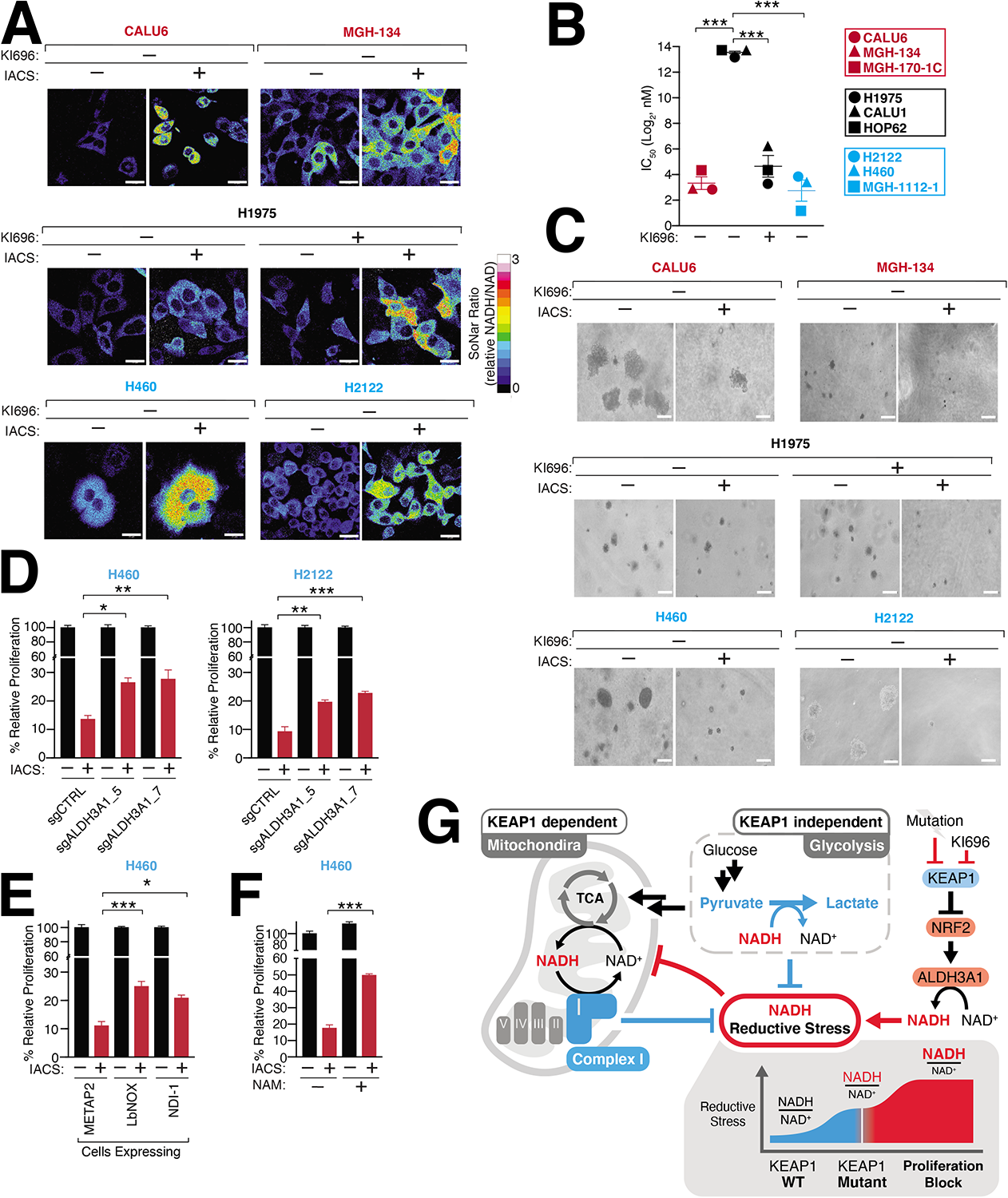
Inducing NADH-reductive stress selectively blocks proliferation of NRF2-activated NSCLCs. (A) IACS-010759 (IACS) a Complex-I inhibitor, selectively increases NADH/NAD^+^ ratio in KEAP1-dependent (red) and KEAP1-mutant cells (blue) but not KEAP1-independent cells. NSCLC cells stably expressing SoNar were treated with IACS and analyzed by immunofluorescence. The representative NADH/NAD^+^ ratiometric image was constructed by taking the ratio of the emission intensity of 408 (NADH binding) vs 488 (NAD^+^ binding) for SONAR (see also Figure S7A). (B) IACS selectively blocks NSCLC proliferation following NRF2 activation. IC_50_ values were calculated for each cell line and where indicated cells were also treated with KI696 (1 µM) (Data are represented as a mean ± SEM, n= 6 biological replicates). (C) IACS selectively inhibits the anchorage-independent growth of NSCLCs with hyperactivate NRF2 signaling. Representative images of NSCLC cell lines grown in soft agar following treatment with IACS-017509 (200 nM) or co-treated with KI696 (1 µM) as indicated (Data are represented as a mean ± SEM, n= 6-8 biological replicates) (see also Figure S7C). (D) Decreasing ALDH3A1 partially rescues IACS-010759 treatment in KEAP1-mutant cells. H460 and H2122 sgRNAs targeting ALDH3A1 and respective controls were treated with IACS and proliferation was determined by crystal violet staining as described in (B) (Data are represented as a mean ± SEM, n= 6 biological replicates). (E-F) Increasing NAD^+^ levels decreases IACS-010759 toxicity. H460 cells expressing DOX-inducible LbNOX, ND1I or METAP2 (control) were treated with DOX (100 nM) for 48 hrs (E) or with 1mM NAM (F) and treated with IACS-010759 (500 nM) and proliferation was determined as described in (B) (Data are represented as a mean ± SEM, n= 6 biological replicates). (G) Model. NRF2 activation following pharmacologic inhibition or mutation of KEAP1 increases ALHD3A1 resulting in NADH reductive stress. KEAP1-dependent and -independent cells utilize different NADH oxidation pathways to counter reductive stress. * indicates *p*-values < 0.05, ** indicates *p*-values < 0.01, *** indicates *p*-values < 0.0001. One-way ANOVA with Sidak’s post-hoc correction was used to determine statistical significance. Scale-bar: 25 μm for immunofluorescence and 50 µM for soft agar.

## Discussion

Activation of NRF2 has been widely regarded as a favorable event in cancer growth. Herein, we provide one explanation for the metabolic cost that a cancer must pay to harbor high NRF2 activity, providing the metabolic context required to support this pathway. By pharmacologically profiling a large panel of genetically diverse NSCLC cell lines in combination with functional genomic analysis, we uncovered, that NRF2 activity can only be tolerated under certain metabolic contexts required to offset high NADH levels. Our findings indicate that NADH levels are both necessary and sufficient for NRF2 sensitivity and that the precise cellular pathways used for NADH oxidation are a key determinant for enduring NRF2 activation. We demonstrate that cells with high glycolytic rates tolerate NRF2 activation and do not incur high levels of NADH reductive stress compared with their counterparts which rely more heavily on Complex I for NADH oxidation. As a result, cells with low glycolytic rates suffer NADH reductive stress imposed by NRF2 activity. Our study highlights a growing appreciation for the role of NADH-reductive stress in cell biology (Goodman et al., 2020; Ho et al., 2017; Perez-Torres et al., 2017; Zhang et al., 2012) and directly demonstrate how high levels of reductive stress mediated through an imbalance in NADH/NAD^+^ directly impacts cancer cell growth. Overcoming oxidative stress has been at the forefront of understanding why cancers activate NRF2. Excitingly, our results illustrate that for a subset of NSCLCs, overcoming NADH reductive stress is a critical determinant for cell proliferation. Cancer dependency (DEPMAP) analysis revealed the essential role of KEAP1 across multiple cancers, suggesting that KEAP1 regulation of NADH-reductive stress may be a general requirement for proliferation. Based on our dissection of this dependency in NSCLCs, we anticipate that many KEAP1-dependent cancers will have lower rates of glycolysis and a high dependence on Complex I for NADH oxidation.

Given the central role of the mitochondria in generating cellular ROS (Murphy, 2009), it is not surprising that NRF2 would modulate mitochondrial activity as one mechanism to downregulate ROS levels. Our study indicates a key role in the manipulation of NADH levels as a mechanism by which NRF2 controls mitochondrial function and supports a growing body of literature connecting NRF2 with mitochondrial regulation (Ding et al., 2021; Sayin et al., 2017). NRF2 launches two parallel campaigns to quelch a rise in oxidative stress by: 1) increasing antioxidant levels and 2) decreasing the cellular sources of oxidative stress, in this case mitochondrial respiration. Our work suggests that NRF2 does not directly modulate mitochondrial gene or protein levels, rather relying on a yet to be described posttranslational mechanism to regulate mitochondria. Given the reductive cellular environment brought upon by NRF2 activation that we describe, it would not be surprising if reductive protein stress brought about by incorrect disulfide bond formation is an additional mechanism by which NRF2 stymies mitochondrial function and cancer growth.

While an abundance of cancer genomics studies indicate that NRF2 activation is common event in NSCLCs, our work begins to illuminate when this activation may be permissible. Loss of KEAP1 is not sufficient to initiate tumor growth and is thought to function in a supportive role following oncogene activation such as KRAS or loss of p53 (DeNicola et al., 2011; Romero et al., 2017; Satoh et al., 2013). Our results suggest that a cancer cell may only be able to support NRF2 activation once its glycolytic rate is increased to overcome the burden of NRF2-induced reductive stress. While loss of NRF2 is not lethal, loss of KEAP1 leads to postnatal lethality due to esophageal hyperkeratinization, disruption of renal function and bone mineralization (Suzuki et al., 2017; Wakabayashi et al., 2003; Yoshida et al., 2018). Whether these dysfunctions arise from NADH reductive stress remains to be determined but underscores that NRF2 activation can only be tolerated in certain metabolic contexts. Undoubtedly, new therapies aimed at activating NRF2 (Robledinos-Anton et al., 2019) in non-glycolytic tissues may have unexpected side effects caused by NRF2-induced reductive stress.

The sensitivity of KEAP1-mutant cells to Complex I inhibition may arise from their limited capacity to oxidize additional NADH. The partial rescue of IACS-010759 following depletion of ALDH3A1 suggests that there are likely additional mechanisms by which NRF2 regulates NADH levels in KEAP1-mutant cells or there are off-targets associated with this inhibitor. While clinical reports suggest that IACS-010759 is tolerable in humans (Lemberg et al., 2022; Yap et al., 2019), its anti-cancer efficacy is limited because of toxicity. While this clinical trial did not rely on biomarkers to guide patient selection our finding suggests IACS-010759 may be useful use in patients harboring KEAP1-mutations.

In summary, our findings provide a complex picture for the role of metabolic tumor suppressors in controlling the proliferation of cancer cells. They provide one explanation for how metabolism-centered oncogenic pathways are supported and the metabolic rewiring required for proliferation upon their activation. Importantly this study identifies a cellular signaling pathway whose activation directly controls reductive stress within cancer cells. Our findings are in line with previous studies demonstrating the critical role that NAD^+^ plays in multiple cellular pathways and how maintenance of NADH homeostasis is required for tumor progression (Birsoy et al., 2015; Diehl et al., 2019; Garcia-Bermudez et al., 2018; Gui et al., 2016; Li et al., 2022; Sullivan et al., 2015). Moreover, the discovery that AMPK, long considered to harbor tumor suppressive functions, is paradoxically required for the growth of tumors by regulating lysosomal gene expression, supports the essentiality of some tumor suppressors (Eichner et al., 2019). Collectively, our study provides a metabolic context by which NRF2 activation can create synthetic lethal opportunities through the generation of NADH reductive stress, forming the basis to exploit this form of stress in treating KEAP1 mutant cancers.

## Acknowledgements

We thank Yi Yang for the SoNar plasmid. We thank all members of the Bar-Peled Lab and David Sabatini for helpful suggestions. This work was supported by grants from the NIH (CA215249, CA260062 to L.B-P, R01AR072304, R01AR043369; P01CA163222; R01CA222871 to D.E.F.), Damon Runyon Cancer Research Foundation Rachleff Innovator Award, AACR (19-20-45-BARP), Paula and Rodger Riney Foundation, the V Foundation, Melanoma Research Alliance, the Mary Kay Foundation and the Ludwig Cancer Center, and the Dr. Miriam and Sheldon G. Adelson Medical Research Foundation.

## Author Contributions

T.W-S. and L.B-P. conceived and designed the study. T.W-S performed most of experiments, with assistance of M.G., A.G., A.C., M.F., H.F. T.V., and V.V., L.S., T-Y.W. performed metabolomic analysis. M.T., M.R., and T-Y. Y. performed proteomics analysis and interpreted the data. C.H.A and T.V analyzed the CRISPR screens with the help of D.E.F. B.R.D generated the model. T.W-S. and L.B-P. wrote the manuscript with assistance from all the coauthors. L.B-P. supervised the studies.

## Competing Interests

L.B-P is a founder, consultant and holds privately held equity in Scorpion Therapeutics. D.E.F. has a financial interest in Soltego, a company developing salt inducible kinase inhibitors for topical skin-darkening treatments that might be used for a broad set of human applications. The interests of L.B-P and D.E.F. were reviewed and are managed by Massachusetts General Hospital and Partners HealthCare in accordance with their conflict-of-interest policies.

## MATERIALS AND METHODS

### KEY RESOURCES TABLE

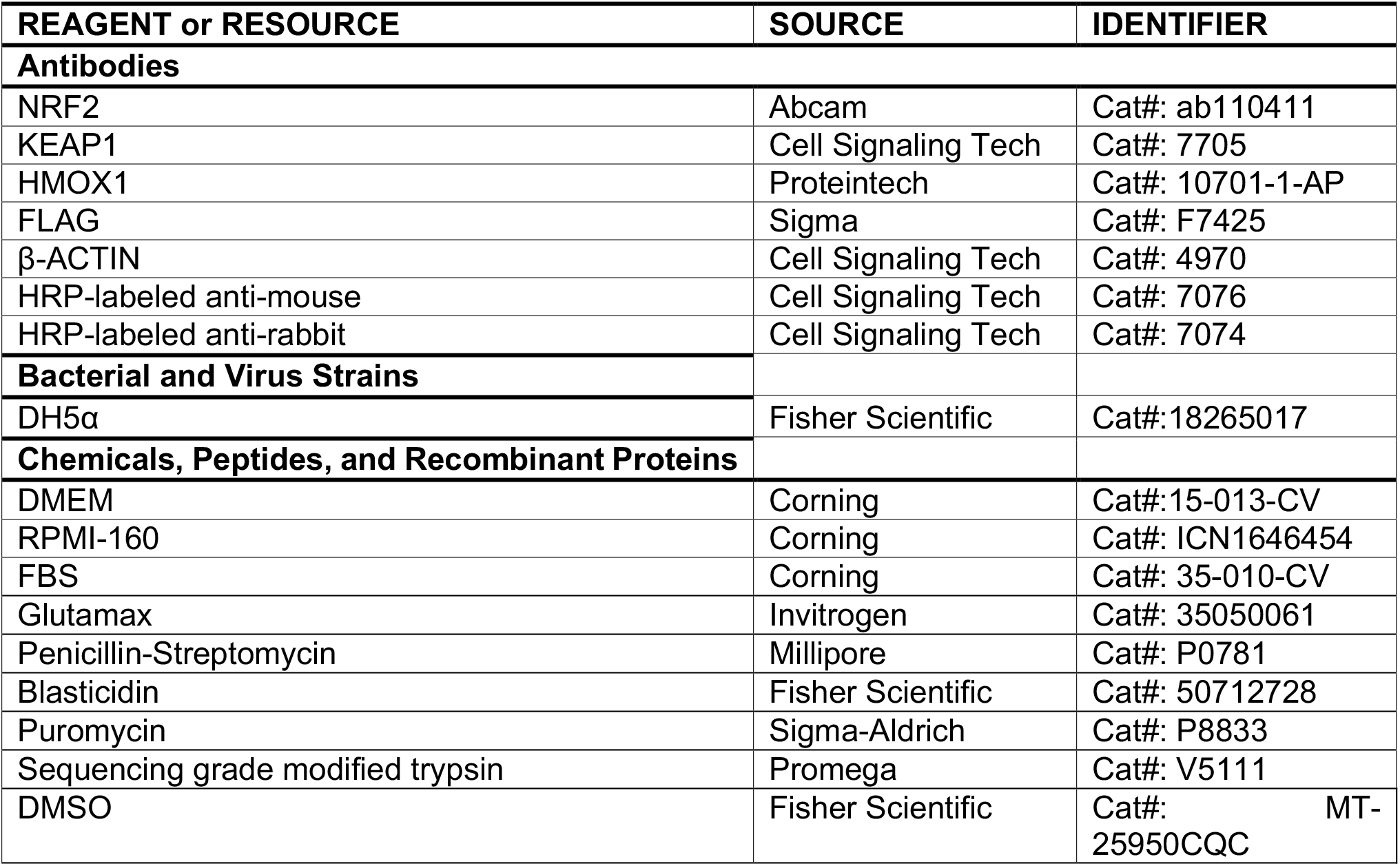
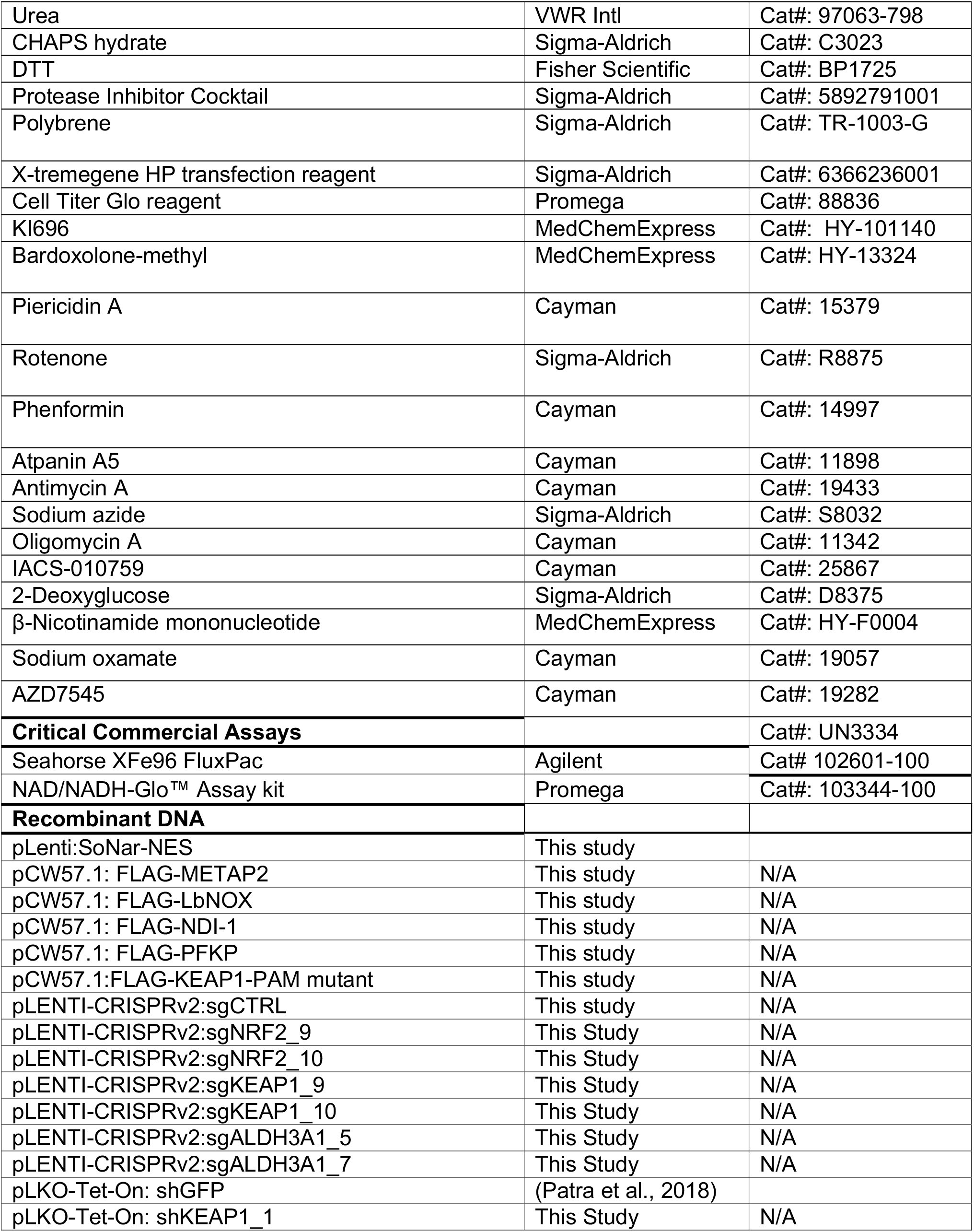
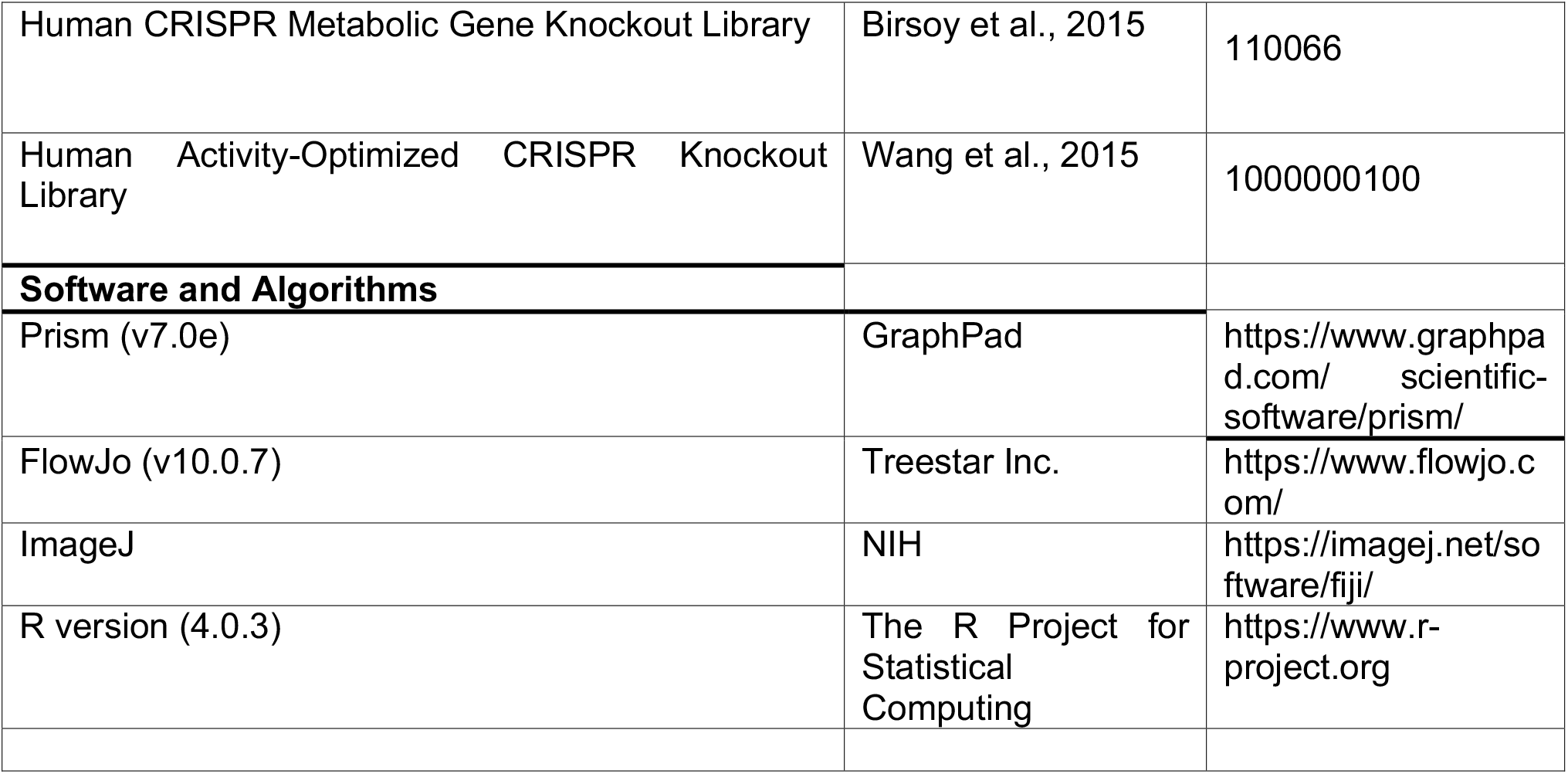

| REAGENT or RESOURCE | SOURCE | IDENTIFIER |
| --- | --- | --- |
| **Antibodies** |  |  |
| NRF2 | Abcam | Cat#: ab110411 |
| KEAP1 | Cell Signaling Tech | Cat#: 7705 |
| HMOX1 | Proteintech | Cat#: 10701-1-AP |
| FLAG | Sigma | Cat#: F7425 |
| β-ACTIN | Cell Signaling Tech | Cat#: 4970 |
| HRP-labeled anti-mouse | Cell Signaling Tech | Cat#: 7076 |
| HRP-labeled anti-rabbit | Cell Signaling Tech | Cat#: 7074 |
| **Bacterial and Virus Strains** |  |  |
| DH5α | Fisher Scientific | Cat#:18265017 |
| **Chemicals, Peptides, and Recombinant Proteins** |  |  |
| DMEM | Corning | Cat#:15-013-CV |
| RPMI-160 | Corning | Cat#: ICN1646454 |
| FBS | Corning | Cat#: 35-010-CV |
| Glutamax | Invitrogen | Cat#: 35050061 |
| Penicillin-Streptomycin | Millipore | Cat#: P0781 |
| Blasticidin | Fisher Scientific | Cat#: 50712728 |
| Puromycin | Sigma-Aldrich | Cat#: P8833 |
| Sequencing grade modified trypsin | Promega | Cat#: V5111 |
| DMSO | Fisher Scientific | Cat#: MT-25950CQC |
| Urea | VWR Intl | Cat#: 97063-798 |
| CHAPS hydrate | Sigma-Aldrich | Cat#: C3023 |
| DTT | Fisher Scientific | Cat#: BP1725 |
| Protease Inhibitor Cocktail | Sigma-Aldrich | Cat#: 5892791001 |
| Polybrene | Sigma-Aldrich | Cat#: TR-1003-G |
| X-tremegene HP transfection reagent | Sigma-Aldrich | Cat#: 6366236001 |
| Cell Titer Glo reagent | Promega | Cat#: 88836 |
| KI696 | MedChemExpress | Cat#: HY-101140 |
| Bardoxolone-methyl | MedChemExpress | Cat#: HY-13324 |
| Piericidin A | Cayman | Cat#: 15379 |
| Rotenone | Sigma-Aldrich | Cat#: R8875 |
| Phenformin | Cayman | Cat#: 14997 |
| Atpanin A5 | Cayman | Cat#: 11898 |
| Antimycin A | Cayman | Cat#: 19433 |
| Sodium azide | Sigma-Aldrich | Cat#: S8032 |
| Oligomycin A | Cayman | Cat#: 11342 |
| IACS-010759 | Cayman | Cat#: 25867 |
| 2-Deoxyglucose | Sigma-Aldrich | Cat#: D8375 |
| β-Nicotinamide mononucleotide | MedChemExpress | Cat#: HY-F0004 |
| Sodium oxamate | Cayman | Cat#: 19057 |
| AZD7545 | Cayman | Cat#: 19282 |
| **Critical Commercial Assays** |  | Cat#: UN3334 |
| Seahorse XFe96 FluxPac | Agilent | Cat# 102601-100 |
| NAD/NADH-Glo™ Assay kit | Promega | Cat#: 103344-100 |
| **Recombinant DNA** |  |  |
| pLenti:SoNar-NES | This study |  |
| pCW57.1: FLAG-METAP2 | This study | N/A |
| pCW57.1: FLAG-LbNOX | This study | N/A |
| pCW57.1: FLAG-NDI-1 | This study | N/A |
| pCW57.1: FLAG-PFKP | This study | N/A |
| pCW57.1:FLAG-KEAP1-PAM mutant | This study | N/A |
| pLENTI-CRISPRv2:sgCTRL | This study | N/A |
| pLENTI-CRISPRv2:sgNRF2_9 | This Study | N/A |
| pLENTI-CRISPRv2:sgNRF2_10 | This Study | N/A |
| pLENTI-CRISPRv2:sgKEAP1_9 | This Study | N/A |
| pLENTI-CRISPRv2:sgKEAP1_10 | This Study | N/A |
| pLENTI-CRISPRv2:sgALDH3A1_5 | This Study | N/A |
| pLENTI-CRISPRv2:sgALDH3A1_7 | This Study | N/A |
| pLKO-Tet-On: shGFP | (Patra et al., 2018) |  |
| pLKO-Tet-On: shKEAP1_1 | This Study | N/A |
| Human CRISPR Metabolic Gene Knockout Library | Birsoy et al., 2015 | 110066 |
| Human Activity-Optimized CRISPR Knockout Library | Wang et al., 2015 | 1000000100 |
| **Software and Algorithms** |  |  |
| Prism (v7.0e) | GraphPad | https://www.graphpad.com/scientific-software/prism/ |
| FlowJo (v10.0.7) | Treestar Inc. | https://www.flowjo.com/ |
| ImageJ | NIH | https://imagej.net/software/fiji/ |
| R version (4.0.3) | The R Project for Statistical Computing | https://www.r-project.org |

### CONTACT FOR REAGENT AND RESOURCE SHARING

Further information and requests for reagents should be directed to the Lead Contact, Liron Bar-Peled.

### EXPERIMENTAL MODEL AND SUBJECT DETAILS

#### Cell lines

All cell lines were grown in RPMI (Corning). Penicillin-Streptomycin (100 mg/ml, Millipore) and 1% GlutaMax (Gibco). All cell lines were tested at least once for Mycoplasma and if not noted elsewhere were obtained from American Tissue Type Collection (ATCC). All *MGH* cell lines were kindly provided by Dr. Aaron N Hata. Whenever thawed, cells were passaged at least three times before being used in experiments. For normoxia and hypoxia experiments, cell lines were maintained at 5% CO_2_ with the indicated oxygen concentrations.

#### Lentivirus virus production

Mammalian lentiviral particles harboring sgRNA-encoding plasmids or cDNA-encoding plasmids were co-transfected with the psPAX2 envelope and VSV-G packaging plasmids into actively growing HEK-293T cells (ATCC) using Xtremegene-HP (Sigma) transfection reagent as previously described (Sarbassov et al., 2005). Virus-containing supernatants were collected 48 hours after transfection, filtered to eliminate cells and target cells were infected in the presence of 8 μg/ml polybrene (Milipore) at a concentration of 2.5 x 10^5^ cells/well. 24 hours later, cells were selected with puromycin (Sigma-Aldrich) or blasticidin (Sigma-Aldrich) and analyzed 3-10 days after selection was initiated. The sequences of sgRNAs and shRNAs used in this study can be found in **Supplemental Table 11**.

#### cDNA cloning and mutagenesis

cDNAs were amplified using Q5 High-Fidelity 2X master mix (NEB) and subcloned into pCW57.1-DOX on or PCW57.1-DOX off (Addgene) or pLentiCRISPRv2 (Addgene) by T4 ligation or Gibson cloning (NEB). Site directed mutants were generated using QuikChange XLII site-directed mutagenesis (Agilent), using primers containing the desired mutations. The sequences of primers used in this study can be found in **Supplemental Table 12.** The SoNar cDNA is described in (Zhao et al., 2015). All constructs were verified by DNA sequencing.

#### Cell lysis

The indicated cell lines were rinsed once with ice-cold PBS and lysed with Triton lysis buffer (1% Triton X-100, 40 mM HEPES pH 7.4, 2.5 mM MgCl_2_ and 1 tablet of EDTA-free protease inhibitor (Roche) (per 25 ml buffer)) and gentle sonication using a QSonica 700A water-chilled sonicator. The soluble fractions of cell lysates were isolated by centrifugation at 13,000 rpm in a microcentrifuge for 10 minutes, normalized and proteins were denatured by the addition of 5X sample buffer and boiling for 5 min as described (Kim et al., 2002). Samples were resolved by 8%–16% SDS-PAGE and analyzed by immunoblotting.

#### Statistical analysis

Statistical analysis was preformed using GraphPad Prism (v10.0.7) for Mac (GraphPad Software) or R statistical programing language (v4.0.3, R-project.org). Statistical values including the exact n, statistical test, and significance are reported in the Figure Legends.

Statistical significance was defined as * p< 0.05 and unless indicated otherwise determined by 2-tailed Student’s t-test or one-way Anova. All post-hoc analyses are indicated in the figure legends. A False Discovery Rate (FDR) was calculated for Proteomics (using Proteome Discover, v2.5, Thermo), RNA-sequencing (EdgeR (Robinson et al., 2010)) and metabolomics (Limma (Ritchie et al., 2015)) analysis to correct for multiple comparisons. CRISPR-scores were calculated as described in CRISPR screen section. For the KI696 and Bardoxolone small molecule screens, a Z-score was calculated using the following method: Z-Score= (x_i_-μ)/σ, where x_i_ is the fold changed (KI696/DMSO) of the i^th^ sample, μ is the mean fold change (KI696/DMSO) across all samples and σ is the standard deviation of fold change (KI696/DMSO) across all samples.

#### Monolayer proliferation assay

Unless otherwise noted, cells were pre-treated with KI696 for 3 days in 6-well plates, prior to the onset of proliferation assays. For cell lines harboring Doxycycline (DOX)-inducible or repressible cDNAs or shRNAs, cells were treated with DOX 3 days in 6-well plates prior to the onset of proliferation assays. For all other compounds, pre-treatment occurred in a 6-well plate for 2 days with the indicated concentrations listed in the figure legends. At the onset of a proliferation assay, cells were cultured in 96-well plates at 2.5 x 10^3^ cells per well in 100 µl of RPMI and the indicated compound or vehicle control was added for compounds where pre-treatment was required. For all other compounds, agents were administered 24 hours after cell seeding. To quantify cell proliferation crystal violet staining was used as described in (Feoktistova et al., 2016)with slight modification. Briefly, culture media was removed and 50 µL of crystal violet stain (0.5% in 25% methanol) was added to cells for 30 min at room temperature. After removal of crystal violet, cells were washed with water and dried overnight before quantification. Cell viability was quantified in ImageJ (NIH v2.0.3) as previously described in (Mehlem et al., 2013) on threshold images. To calculate half maximal inhibitory concentrations (IC_50_) cells were cultured at 2.5 x 10^3^ cells per well in 100 µL RPMI media and compounds were added the following day. Cell viability was assessed on day six of treatment using crystal violet staining. IC_50_ values were calculated using log(inhibitor) vs % normalized response formula in Prism v.7 (GraphPad). For Figure S1I, cells transduced with the indicated sgRNAs were seeded in a 96-well plate in 100 µL of media and 50 µl of Cell Titer Glo reagent (Promega) was added to each well and the luminescence read on the SpectraMax M5 plate reader (Molecular Devices).

#### Anchorage-independent growth assay

Multiple NSCLC cell lines were tested for their ability to form colonies in soft agar. Cells were seeded at concentrations between 3.0-6.0 x 10^4^ cells/well in a 6-well plate, cell concentrations required to form viable colonies in a 3 week time frame. CALU6, H460 and H2122 were seeded at 3.0 x10^4^ per well. MGH-134 and H1975 were seeded at 6.0×10^4^ cells/well. Where indicated, cells were pre-treated with 1 µM of KI696 for 3 days prior to the onset of anchorage independent growth. For all assays, an equal number of cells from each comparison group was embedded in a solution of 0.4% Noble agar solution (Difco Labs) and the indicated agents or vehicle control were added to the 0.4% Noble agar solution before solidification. Cells were then placed on top of hardened layer of 0.6% agar in a 6-well plate. Cells were grown for 14-20 days at 37°C with 5% CO_2_. Fresh media (200 µL) was added every 5 days. Bright field images were obtain using light microscopy (Nikon Eclipse Ti) with x10 objective lens and colony formation area was measured in ImageJ (NIH, v2.0.3) as described in (Mehlem et al., 2013).

#### Genome-wide and metabolism focused CRISPR screens

CRISPR screens were conducted as previously described in (Wang et al., 2015). Briefly, CALU6 cells were infected with a genome-wide CRISPR library (Addgene, (Wang et al., 2015)) or CALU6 and H1975 cells were infected with a metabolism focused CRISPR library (Addgene,(Birsoy et al., 2015)) ensuring a multiplicity of infection ∼0.3 following 3 days puromycin selection. Cells were allowed to recover for 1 day and an initial input was taken with the number of infected cells corresponding to 1000X the size of the library. The screen was initiated by treating cells (∼200×10^6^ cells for genome-wide and 30×10^6^ for metabolism-focused) with DMSO or 1 µM KI696, maintaining this cell number and compound for 11 population doublings. At the end of the screen, cells were harvested, and genomic DNA was extracted using Macherey Nagel Blood XL kit (Macherey). Libraries were generated from each sample by PCR based amplification of the sgRNA amplicon from 200 µg of genomic DNA using custom PCR primers harboring an index primer and illumina 5’ and 3’ adaptors. Libraries were pooled and analyzed on a NextSeq500 (Illumina) use single end 75bp reads. sgRNAs were mapped and quantified as describe in (Wang et al., 2015). The enrichment for each sgRNA was calculated by taking the Log_2_ ratio of (sgRNA counts, treatment/sgRNA counts input). The CRISPR score for each gene was calculated by subtracting the enrichment score for KI696 treated samples from the enrichment score for DMSO treated samples.

#### Gene set enrichment analysis (GSEA) and metabolic pathway enrichment

GSEA (Subramanian et al., 2005)was carried out using pre-ranked lists from genome-wide CRISPR-score values using the fast GSEA package in (Korotkevich et al., 2021) Gene sets were collected from MSigDB version 7.4 and top/bottom 250 gene categories were then stratified into subcategories as described in Figure 1F. To identify KI696 mediated sensitivity from different metabolic pathways, genes were curated into different metabolic pathways as described in (Levy et al., 2016) and a CRISPR value for each pathway was determined by calculating the aggregate CRISPR-score for each gene comprising this pathway (see Figures 1H, S2A).

#### Glycolytic signature

A glycolytic gene signature was calculated for NSCLC cell lines according to (Haynes et al., 2017) with slight modifications. Briefly, the mean expression of glycolytic genes, as defined in KEGG glycolysis gene set V7.5.1, was calculated for NSCLC cell lines by using publicly available transcriptomic data (20Q4) from DEPMAP (Ghandi et al., 2019). Expression values were then Z-score normalized to get glycolytic gene-score = (x_i_-μ)/σ, where x_i_ is the mean expression value of glycolytic genes of the i^th^ sample (NSCLC cell line), μ is the average expression of glycolytic gene across all samples and σ is the standard deviation of glycolytic gene expression across all samples.

#### Seahorse Flux Analyses

Oxygen consumption rates (OCR) and extracellular acidification rates (ECAR) were measured using XFe96 Extracellular Flux Analyzer (Agilent) as described previously with slight modifications^1^. Briefly, all cell lines were plated on a poly-L-lysine coated 96-well Seahorse plates (Agilent) at 10 x 10^3^ cells/well, with the exception of H522 and H661 that were seeded at 15 x 10^3^ and 5 x 10^3^ cells/well, respectively. Cells were treated with KI696 (1 μM) or DMSO for 48 hours in RPMI media. To analyze OCR and ECAR, the media was changed to RPMI supplemented with 2 mM L-glutamine and 10 mM D-glucose, but lacking sodium bicarbonate. Measurements of OCR were carried out at baseline and after injections of oligomycin (1.5 μM), FCCP/Na Pyruvate (3 μM/1 mM), and Antimycin (1 μM). To measure ECAR, the assay media was modified to RPMI supplemented with 2 mM L-glutamine and ECAR was measured following the injections of D-glucose (10mM), oligomycin (1.5 μM), and 2-Deoxyglucose (100mM). OCR/ECAR measurements were carried out at baseline. All OCR and ECAR values were normalized to total protein content as measured by BCA (Pierce) according to manufacturer’s instructions.

#### Confocal imaging of cell lines expressing SoNar reporter

NSCLC cell lines expressing the indicated the SoNar reporter were seeded on poly-lysine coated 8-well chamber (iBidi) at 10×10^3^ cells per well and treated with compounds as described in the text. For KI696 treatments, cells were pre-treated for 2 days with 1 µM KI696 or vehicle control prior to seeding on glass bottom dishes. Dishes were firmly mounted the stage adaptor of Zeiss 710 Laser Scanning Confocal microscope (Carl Zeiss Inc.). Constant temperature (37°C), humidity, and 5% CO_2_ atmosphere are maintained throughout the duration of cell imaging. Images were acquired using a 63X oil objective. Relative NAD^+^ levels were determined by exciting SoNar expressing cells with a 488-nm laser and measuring emission at 500-520 nm range. Relative NADH levels were determined by exciting SoNar expressing cells with a 405-nm laser and measuring emission at 500-545 nm range. Acquisition parameters were kept identical between samples. Images were acquired using a 63X oil objective. Ratiometric images of SONAR were processed using ImageJ (NIH, v2.0.3) to 32-bit images and presented in 16 colors mode. Threshold images were quantified for mean fluorescence intensity in ImageJ.

#### Flow cytometry analysis

NSCLC cells expressing the indicated SoNar reporter were seeded at 0.25 x 10^6^ cells/well in a 6-well plate for 48 hours. Cells were dislodged by trypsin digestion, resuspended in FACS buffer (PBS+1% FBS, 2mM EDTA) and treated sequentially at room temperature with 0.5 µM Rotenone (Sigma-Aldrich) followed by 5mM sodium oxamate (Sigma-Aldrich). The fluorescent signal at 530 nm following excitation at 405 nm (NADH binding) or 488 nm (NAD binding) was measured every 2 min for a total of 30 min by an Aurora (Cytek) flow cytometer. The ratio of λex = 488 nm/ λem = 530 nm to λex = 405 nm/ λem = 530 nm signal was determined using Flowjo v10.6.

#### Lactate dehydrogenase (LDH) activity

LDH activity was determined as described in (Zdralevic et al., 2018). Briefly, cell pellets were lysed in water and the supernatant was cleared by centrifugation at 13,000 x rpm for 10 min at 4°C. Protein concentration was determined by Bradford (Bio-Rad). To initiate the NADH oxidation assay equal concentrations of cell lysate (20 µL) were added into 80 µL of a reaction buffer (1mM sodium pyruvate, 0.5 mM NADH and 200 mM Tris pH 7.5). NADH oxidation rate was determined by kinetic absorbance at 340 nm which was collected every 2 min for 30 min using SpectraMax M5 plate reader (Molecular Devices). To determine LDH activity, NADH half-life was determined by one-phase exponential decay model using Prism v7 (GraphPad) and LDH activity is represented as 1/NADH_(half-life)_.

#### Lactate measurement

Cell lines were cultured at 1×10^6^ cells per well in a 6-well plate in 2mL of RPMI culture media. The following day, culture media was replaced with 1 mL of serum-free RPMI after three successive washes with PBS. After 8 hours, the entirety of the supernatant (1 mL) was collected and mixed with a 50 μL of reaction buffer (30 mM Tris (Sigma-Aldrich) pH 7.4, 40 μM resazurin (Sigma-Aldrich), 0.01U rLDH (Abcam), 0.05U Diaphorase (Sigma-Aldrich), 1mM NAD^+^ (Cayman). Reactions were carried out at room temp for 30 min in an opaque 96-well plate. The fluorescent signal corresponding to NADH oxidation was measured (550-585nm) using SpectraMax M5 plate reader (Molecular Devices) and normalized to cell number.

#### GCMS analysis

1×10^6^ cells were cultured in a 6 cm dish in a total of 5mL RPMI media and treated with DMSO or KI696 (1μM) for 48 hours. Cells were washed with 0.9% NaCl solution and immediately flash frozen in liquid nitrogen. Metabolite extraction was undertaken by adding 800 µL ES1 (H_2_O:L-Norvalin (1mg/mL):Glutarate (1mg/mL), 15mL:37.5μL:37.5μL) (all from Sigma-Aldrich) followed by addition of chloroform (500μL). Samples were vortexed, centrifuged at 13,000, x rpm for 10 min, 4°C and the upper phase was dried and derivatized with TBDMS (Sigma-Aldrich) for downstream GC-MS analyses. Samples were analyzed on an Agilent 7890B GC system was coupled to 5977B single quadrupole mass spectrometer equipped with an electron ionization source. Automated injections were performed with an Agilent 7693 autosampler. The injector temperature was held constant at 270 °C. Injections of 1 μL were made in splitless mode. Chromatography was performed on a HP-5ms Ultra Inert Column (30 m x 0.25 mm, 0.25 μm film thickness, Agilent). Helium carrier gas was used at a constant flow of 1ml/min. The GC oven temperature program was 100 °C initial temperature with 3 min hold time and ramping at 10 °C/min to a final temperature of 300 °C with 12 min hold time. The transfer line temperature was 250 °C, and the source temperature was 230 °C. After a solvent delay of 5.5 min, mass spectra were acquired at 2.9 scans/s with a mass range of 50 to 550 m/z. Data processing was performed with MassHunter Workstation Software Quantitative Analysis (Version B.09.00 / Build 9.0.647.0, for GCMS and LCMS).

#### LCMS analysis

2.5×10^5^ cells were cultured in a 6 well plate in 4mL of RPMI and treated with DMSO or KI696 (1μM) for 48 hours. On the day of sample collection, cells were washed three times with a 75 mM ammonium carbonate solution followed by extraction with 70% ethanol at 70°C for 3 minutes. The supernatant was cleared by centrifugation at 13,000 x rpm for 10 min at 4°C and immediately stored in −80°C until further processing by LCMS.

LCMS analysis was performed on a platform consisting of an Agilent 1260 Infinity II LC pump coupled to a Gerstel MPS autosampler (CTC Analytics, Zwingen, Switzerland) and an Agilent 6550 Series Quadrupole TOF mass spectrometer (Agilent, Santa Clara, CA, USA) with Dual AJS ESI source operating in negative mode as described previously (Fuhrer et al, 2011). The flow rate was 150 µl/min of mobile phase consisting of isopropanol:water (60:40, v/v) with 1 mM ammonium fluoride. For online mass axis correction, two ions in Agilent’s ESI-L Low Concentration Tuning Mix (G1969-85000) were used. Mass spectra were recorded in profile mode from m/z 50 to 1,050 with a frequency of 1.4 s for 2 × 0.48 min (double injection) using the highest resolving power (4 GHz HiRes).

All steps of mass spectrometry data processing and analysis were performed with MATLAB (The Mathworks, Natick, MA, USA) using functions embedded in the Bioinformatics, Statistics, Database, and Parallel Computing toolboxes as described previously in (Fuhrer et al., 2011). The resulting data included the intensity of each mass peak in each analyzed sample. Peak picking was done for each sample once on the total profile spectrum obtained by summing all single scans recorded over time, and using wavelet decomposition as provided by the Bioinformatics toolbox. In this procedure, a cutoff was applied to filter peaks of less than 5,000 ion counts (in the summed spectrum) to avoid detection of features that are too low to deliver meaningful insights. Centroid lists from samples were then merged to a single matrix by binning the accurate centroid masses within the tolerance given by the instrument resolution. Starting from the HMDB v4.0 database (Wishart et al., 2007), we generated a list of expected ions including deprotonated, fluorinated, and all major adducts found under these conditions. All formulas matching the measured mass within a mass tolerance of 0.001 Da were enumerated. As this method does not employ chromatographic separation or in-depth MS2 characterization, it is not possible to distinguish between compounds with identical molecular formula. The confidence of annotation reflects Level 4 but – in practice - in the case of intermediates of primary metabolism it is higher because they are the most abundant metabolites in cells biological extracts. The resulting matrix lists the intensity of each mass peak in each analyzed sample. An accurate common m/z was recalculated with a weighted average of the values obtained from independent centroiding.

#### RNAseq analysis

CALU6 cells were cultured in 6-well plates (2.5 x 10^5^ cells/well) and treated with KI696 (1 µM) or vehicle control for 48 hrs. RNA was isolated by RNeasy Kit (Qiagen) following the manufacturer’s instructions and digested with DNase (Qiagen) from n=2 samples per condition. mRNA-seq libraries were prepared using NEBNext® Poly(A) mRNA Magnetic Isolation Module (NEB) kit according to manufacturer’s instructions. Libraries were then quantified by Kappa Library Quantification (Roche), pooled, and sequenced by single-end 75 base pairs using the Illumina NextSeq 500 platform. FASTQ files were then processed using RNA Express module (Illumina) and raw counts were further processed in R (v.4.0.3) and edgeR (PMID: 19910308) to obtain relative gene expression.

#### Proteome wide analysis of NRF2 activation

CALU6 cells cultured in 6-well plates (0.25 x 10^6^ cells) and treated with 1 µM KI696 for 48 hrs. Cells were washed once with ice-cold PBS, snap-frozen in liquid nitrogen and stored at −80°C until use. Frozen cell pellets were lysed in PBS supplemented with Benzonase (Santacruz) and protease inhibitors (Roche) using a chilled bath sonicator (QSONICA) and centrifuged for 3 min at 300 *g*. Proteins were quantified by BCA assay (Thermo Scientific) and a total of 50 μg of total protein extracts were used for each TMT channel. Protein extracts were reduced with 5 mM 5-Tris (2-carboxyethyl) phosphine hydrochloride (TCEP) (Sigma-Aldrich) for 2 min at room temperature, followed by alkylation using 20 mM Iodoacetamide (Sigma-Aldrich) for 30 min in the dark at room temperature. SP3 magnetic beads (Cytiva) were prewashed with LC-MS grade water and 250 µg combined SP3 beads (1:1 hydrophobic:hydrophilic) and LC-MS grade ethanol were added to each sample to reach a final concentration of 50% ethanol. SP3 protein binding occurred for 30 min at room temperature and beads were subsequently washed 3 times with 80% ethanol for resuspended with 175 ul of Trypsin/Lys-C (1 µg, Thermo Scientific A40009) in 200 mM EPPS (pH 8.4)/5 mM CaCl_2_. Proteins were digested overnight (16 h) at 37°C and peptides were dried using a Speedvac (Thermo Scientific). Peptides were desalted with stage tips using the following procedure: Peptides were reconstituted with 5% acetonitrile/0.1 % formic acid and loaded onto Empore C18 disks (3M) packed into a 200 µl pipette tips pre-equilibrated with LC-MS grade methanol and water containing 0.1% formic acid. C18 disk were washed 10 times with LC-MS grade water containing 0.1 % formic acid and subsequently eluted with 80% acetonitrile/0.1% formic acid and dried using a Speedvac (Thermo Scientific). Peptides were quantified with Quantitative Colorimetric Peptide Assay (Thermo Scientific) and 5 µg of total peptides were labeled using TMTpro16-plex reagents (Thermo Scientific). Briefly, peptides were reconstituted with 30% acetonitrile/70% 200 mM EPPS (pH 8.4) and labeled with 50 µg of TMT reagent per channel for 75 min at room temperature with rotation. Labeling was terminated by the addition of 5% hydroxylamine (Acros Organics) for 15 min, followed by supplementation with 10% formic acid. Samples were pooled and dried using a Speedvac (Thermo) and fractionated using high pH reversed phase fractionation (Thermo Scientific) following the manufactures directions.

Fractionated samples were reconstituted with 2% acetonitrile/0.1 % formic acid and subjected to liquid chromatography using an Easy nLC 1200. The columns used are an Acclaim PepMap trap Column (75 μm x 20 mm packed with 3-μm particles of C18 stationary phase) and an EASY-Spray analytical column (75 μm x 500 mm packed with 2-μm particles of C18 stationary phase). Peptides were loaded onto the trap column using 0.1 % formic acid in water. Peptides were separated over a 190 min gradient, consisting of 5-28% solvent B over 155 min, 28-43% solvent B over 25 min, 43-95% solvent B over 10 min, and 90% solvent B held for 10 min. Solvent A was 0.1% formic acid in water and solvent B was 80% acetonitrile and 0.1% formic acid in water. Peptides were ionized at 2300 V and analyzed on an Orbitrap Eclipse mass spectrometer coupled with a FAIMSpro system. Ionized peptides were separated using FAIMSpro (1.5 sec per cycle). MS1 was analyzed in the orbitrap at 120,000 resolution using 50 ms ion injection time (IIT) and 250% automatic gain control (AGC). Collision energy was set to 36% (CID) and fragmented ions were detected in the ion trap using default IIT and AGC. Real time search function was enabled to detect human peptides. Synchronous precursor selection was enabled and was set to isolate twenty notches. MS3 was analyzed in the orbitrap with the mass range at 100 – 500 Da and the resolution at 50,000.

Acquired RAW data were analyzed using ProteomeDiscoverer v2.5 (Thermo Scientific) using the SequestHT module. Data was search against the UniProtKB human universal database (UniProt UP000005640, downloaded May 2020) combined with the common Repository of Adventitious Proteins (cRAP, classes 1,2,3,and 5). Parameters were set as follows: MS1 tolerance of 10 ppm, MS/MS mass tolerance of 0.6 Da, trypsin (full) digestion with a maximum of zero missed cleavages, minimum peptide length of 6 and maximum of 144 amino acids. Cysteine carbamidomethylation (57.021 Da) and methionine oxidation (15.995 Da) were set as dynamic modifications. Lysine- and N-ternimous-TMTpro modification (304.207 Da) was set as static modification. A false discovery rate (FDR) of 1% was set for peptide-to-spectrum matches using the Percolator algorithm and for protein assignment. Reporter ion quantification was based on signal over noise (S/N) with the co-Isolation Threshold at 50, S/N threshold at 10, and SPS mass matches threshold at 50%. Abundances was normalized to total peptides. Protein ratio was calculated using PD2.5 pairwise ratio based algorithm and t-test was used for significance.

#### NADH/NAD measurement

0.25×10^6^ CALU6 cells were treated with 1 µM KI696 or vehicle control in 6-well plates for 48 days. The ratio NADH/NAD^+^ was determined NAD/NADH-Glo™ Assay Kit (Promega) following the manufacturer’s protocol. Absorbance was measured using a SpectraMax M5 plate reader (Molecular Devices).

---

**Figure S1:**
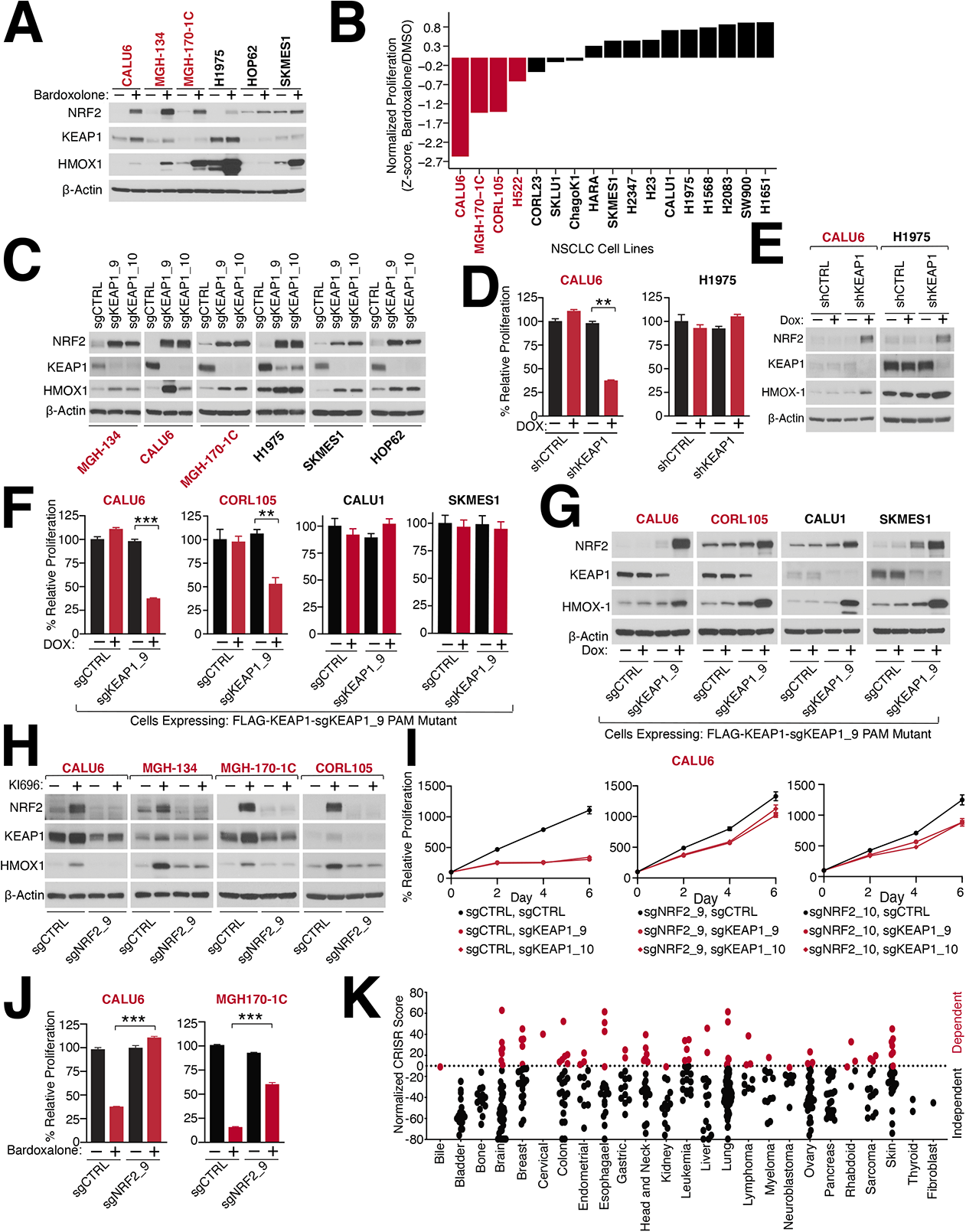
Identification of KEAP1-mutant NSCLC cell lines. (A) Bardoxolone treatment activates NRF2. Representative immunoblot analysis of NSCLC cell lines treated with bardoxolone (100 nM) for 48 hrs. (B) Bardoxolone blocks the proliferation of KEAP1-dependent cells. NSCLC cells were treated with bardoxolone (100 nM) and proliferation was determined by crystal violet staining 6 days post treatment. (C) Representative immunoblot analysis of NSCLC cell lines expressing the indicated sgRNAs targeting KEAP1. (D) shRNA-mediated depletion of KEAP1 blocks the proliferation of KEAP1-mutant cells. CALU6 and H1975 were infected with a doxycycline (DOX)-inducible shRNA targeting KEAP1 or a control. Cells were pre-treated with DOX (100 nM) for 72 hrs and cell proliferation was determined as described in (B) (Data are represented as a mean ± SEM, n= 3-5 biological replicates). (E) Representative immunoblot analysis of DOX-inducible depletion of KEAP1. (F-G) Addback of sgRNA resistant KEAP1 cDNA rescues KEAP1 depletion. NSCLC cell lines stably co-expressing a DOX-repressible FLAG-KEAP1-PAM mutant cDNA and sgRNAs targeting KEAP1 were treated with DOX (100 nM) for 72 hrs and cell proliferation (F) was determined as described in (B) and levels of the indicated proteins were determined by immunoblot (G) (Data for proliferation assay are represented as a mean ± SEM, n= 3-5 biological replicates). (H) Representative immunoblot analysis of NSCLC cell lines expressing the indicated sgRNAs targeting NRF2 following treatment with KI696 (1 µM). (I) Depletion of NRF2 rescues the proliferation of CALU6 cells following loss of KEAP1. The proliferation of CALU6 cells co-expressing sgRNAs targeting NRF2 or KEAP1 was assessed by measuring relative concentrations of ATP (Data are represented as a mean ± SEM, n=6 biological replicates). (J) NRF2-depletion rescues bardoxolone sensitivity in KEAP1-dependent cell lines. NSCLC cells expressing the indicated sgRNA targeting NRF2 or a control were treated with Bardoxolone and proliferation was determined as described in (B) (Data are represented as a mean ± SEM, n= 5 biological replicates). (K) KEAP1 is a dependency in multiple cancer subtypes as indicated in red. Cancer dependency (DEPMAP(Tsherniak et al., 2017)) data was analyzed as described in (Lenoir et al., 2018). * indicates *p*-values < 0.05, ** indicates *p*-values < 0.01, *** indicates *p*-values < 0.0001. One-way ANOVA with Sidak’s post-hoc analyses used to determine statistical significance.

**Figure S2:**
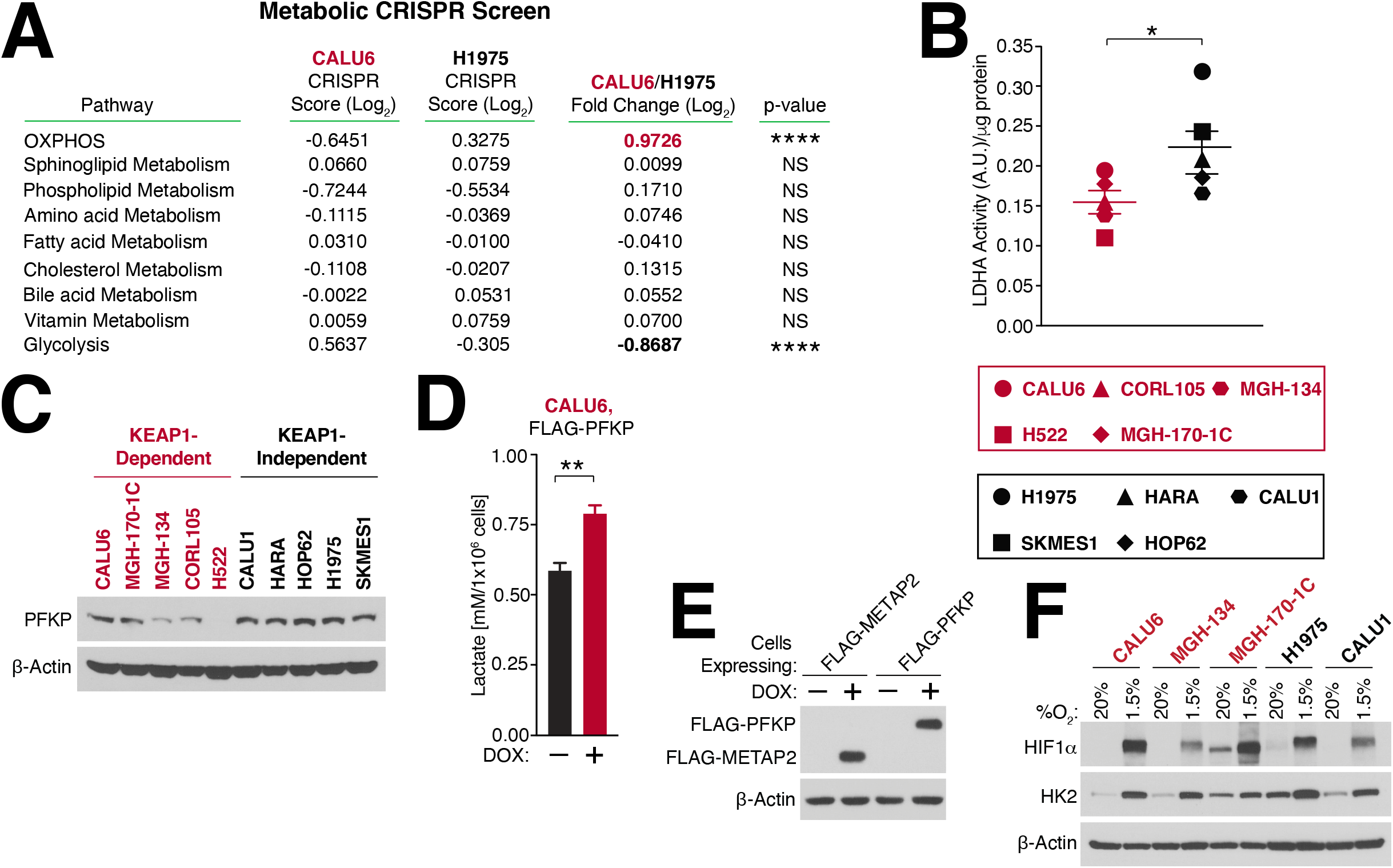
KEAP1-independent cells are marked by high glycolytic rates. (A) Table summarizing CRISPR scores for the indicated metabolic pathways in CALU6 and H1975 cells (see also Figure 1H). (B) KEAP1-dependent cells have lower lactate dehydrogenase (LDH) activity in comparison to KEAP1-independent cells. LDH activity was determined for each cell line as described in the methods (Data are represented as a mean ± SEM, n= 5 samples per group with 16 measurements per sample). (C) PFKP is differentially expressed in KEAP1-independent cell lines. Representative immunoblot analysis of the indicated proteins in KEAP1-dependent and KEAP1-independent NSCLC cell lines (see also Figure 2C). (**D**) Over-expression of PFKP increases lactate levels in CALU6 cells. CALU6 cells stably expressing DOX-inducible FLAG-PFKP were treated with DOX (100 nM) for 3 days and lactate levels were measured as described in the methods (Data are represented as a mean ± SEM, n= 3 biological replicates). (E) Immunoblot analysis of FLAG-PFKP or FLAG-METAP2 expression in CALU6 cells following DOX treatment (100 nM). (F) Induction of hypoxia in NSCLC cell lines. NSCLC cell lines were grown at the indicated oxygen concentrations for 3 days and the expression of the indicated proteins was determined by immunoblot. * indicates *p*-values < 0.05, ** indicates *p*-values < 0.01, *** indicates *p*-values < 0.0001. Statistical significance was determined by Student’s t-test.

**Figure S3:**
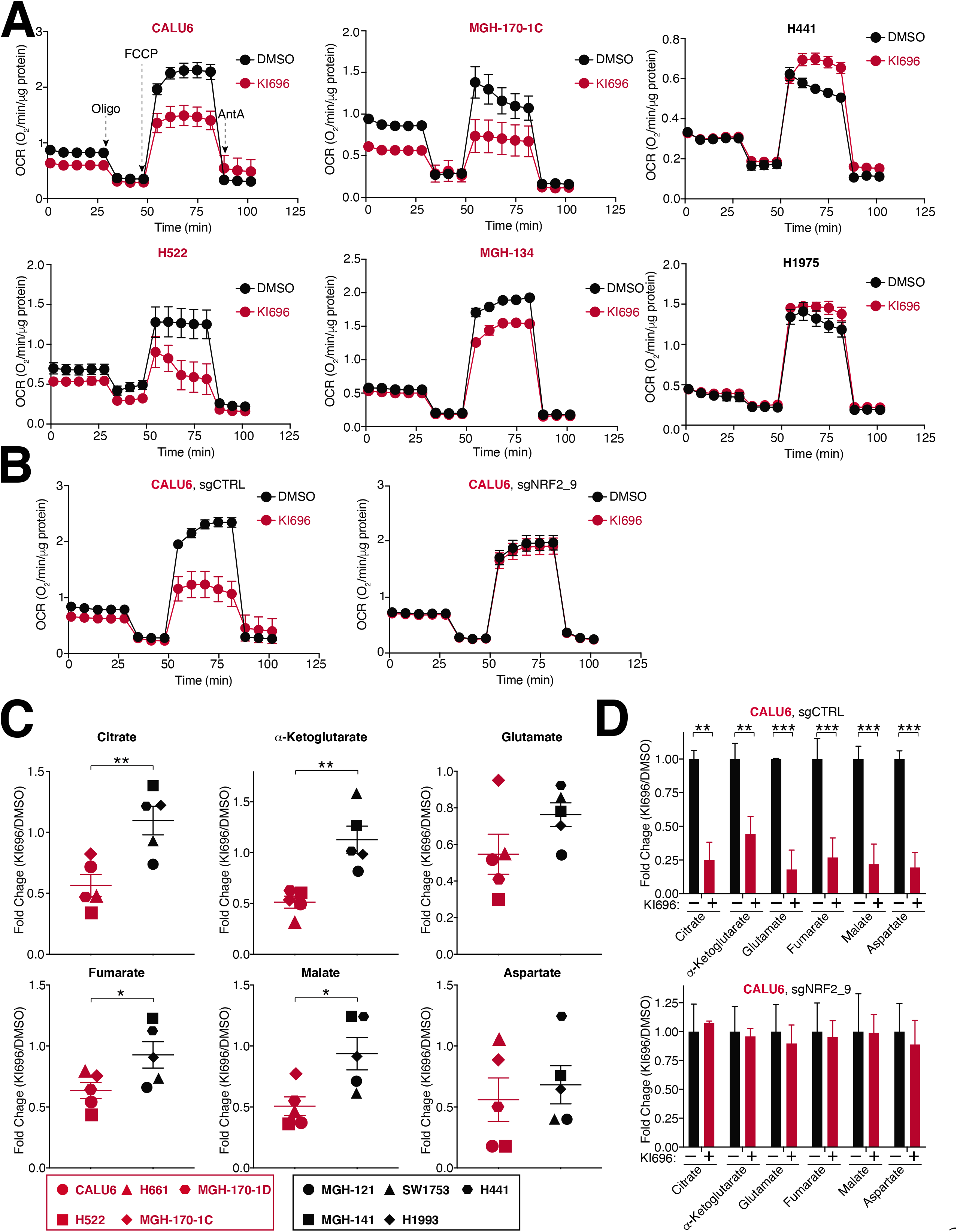
NRF2 activation decreases mitochondrial activity in KEAP1-dependent cells. (A) NSCLC cell lines were treated with KI696 (1 µM) for 48 hrs and the oxygen consumption rate (OCR) was determined using a Seahorse Bioflux analyzer (Data are represented as a mean ± SEM, n= 6-8 biological replicates). (B) NRF2 regulates OCR in KEAP1-dependent cells. CALU6 cells expressing the indicated sgRNA targeting NRF2 or a control, were treated with KI696 (1 µM) for 48 hrs and OCR was determined as described in (A) (Data are represented as a mean ± SEM, n= 6-8 biological replicates). (C) NRF2 activation decreases TCA metabolites in KEAP1-dependent cell lines. NSCLC cell lines were treated with KI696 (1 µM) for 48 hrs and the levels of the indicated metabolites were determined by GCMS (see methods). Fold change (KI696/DMSO) is depicted in the plots (Data are represented as a mean, n= 5 samples per group with 4 biological replicates per sample). (D) NRF2 regulates TCA metabolism in KEAP1-dependent cells. CALU6 cells expressing the indicated sgRNA targeting NRF2 or a non-targeting control, were treated with KI696 and the levels of the indicated metabolites were determined by LCMS (see methods). Fold change (KI696/DMSO) is depicted in the plots (Data are represented as a mean ± SEM, n=3 biological replicates). * indicates *p*-values < 0.05, ** indicates *p*-values < 0.01, *** indicates *p*-values < 0.0001. Statistical significance was determined by Student’s t-test and corrected for multiple hypotheses (D) by False Discovery Rate (FDR), see Methods.

**Figure S4:**
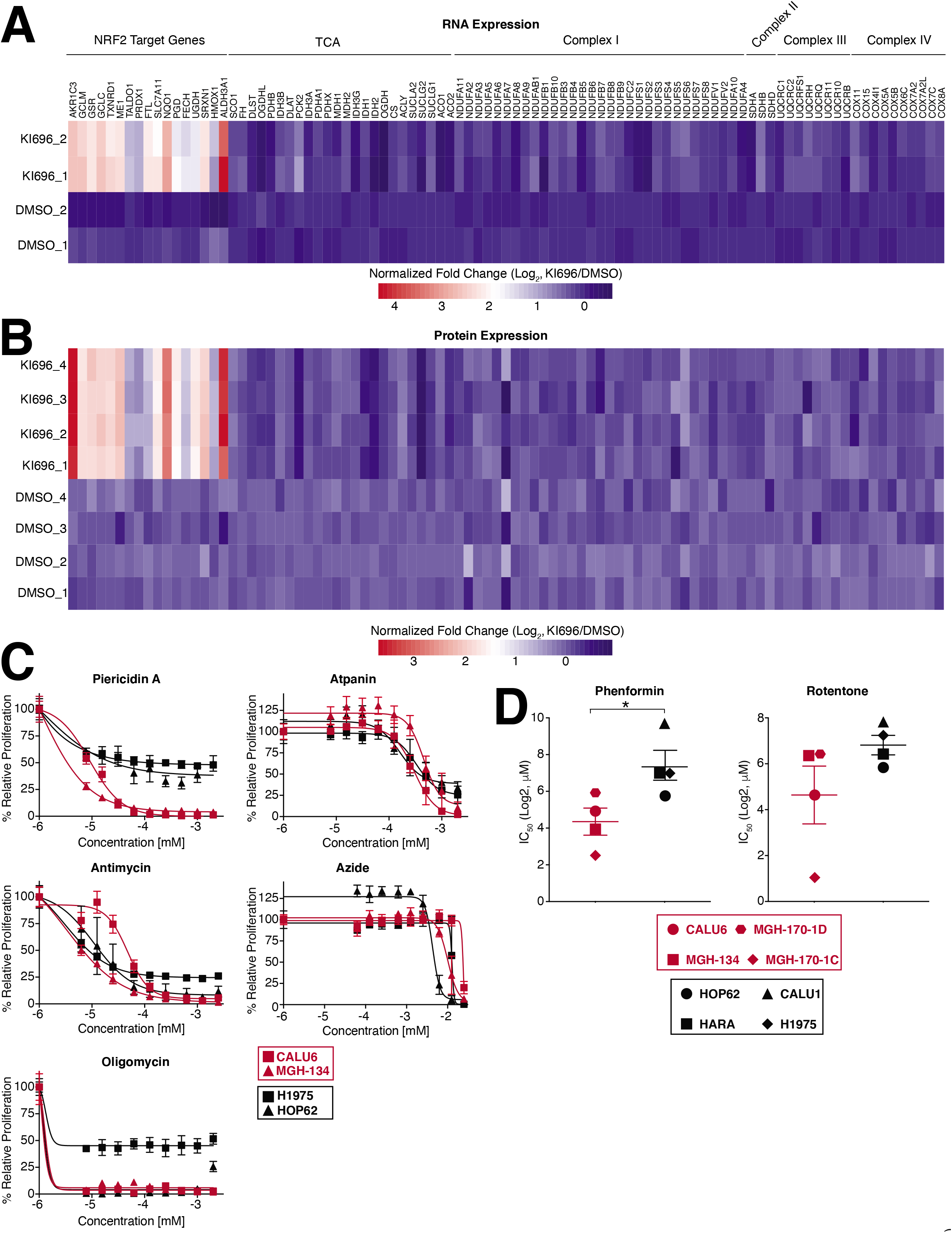
KEAP1-dependent cells are sensitive to Complex I inhibition. (A-B) NRF2 activation does not change the expression of mitochondrial genes involved in ETC or TCA cycle. CALU6 cells were treated with KI696 (1 µM) for 48 hrs and changes in gene expression (A) were determined by RNAseq (see methods and **Supplementary Table 8**) or protein expression (B) were determined by proteomics and depicted in the corresponding heatmaps (see methods and **Supplementary Table 10**). (C) Proliferation of NSCLC cell lines following treatment with the indicated ETC inhibitors following 6 days of treatment were determined by crystal violet staining (Data are represented as a mean ± SEM, n= 6 biological replicates). (D) Complex I inhibitors selectively block KEAP1-dependent cell lines. IC_50_-values were determined for a panel of NSCLC cell lines (Data are represented as a mean ± SEM, n= 4 samples per group and 6 biological replicates). * indicates *p*-values < 0.05, ** indicates *p*-values < 0.01, *** indicates *p*-values < 0.0001. Statistical significance was determined by Student’s t-test.

**Supplementary Figure 5:**
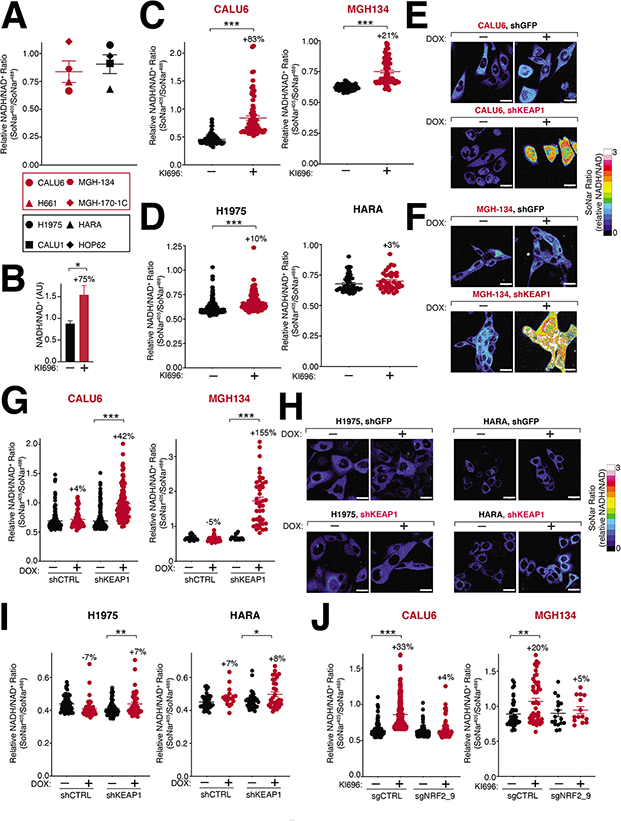
NRF2 regulates the NADH/NAD^+^ ratio in KEAP1-dependent cells. (A) KEAP1-dependent and KEAP1-independent cells have a similar NADH/NAD^+^ ratio at basal states. The NADH/NAD^+^ ratio in NSCLC cell lines stably expressing SONAR was determined by flow cytometry with the ratio of the emission intensity at λem 530nm after excitation λex at 408nm (NADH binding) compared to excitation λex at 408nm 488 (NAD^+^ binding) (Data are represented as a mean ± SEM, n= 4 samples per group). (B) NRF2 activation increases the NADH/NAD^+^ ratio in CALU6 cells. CALU6 was treated for 2 days with KI696 (1 µM) and the NADH/NAD^+^ ratio was determined using a coupled enzymatic method as described in methods (Data are represented as a mean ± SEM, n= 3 biological replicates). (C) Quantification of NADH/NAD^+^ ratio following NRF2 activation in KEAP1-dependent cells. NSCLC cell lines expressing SONAR were treated for 2 days with KI696 and the NADH/NAD^+^ ratio was determined by immunofluorescence analysis comparing the emission intensity after excitation at λex 408nm to λex 488nm (Data are represented as a mean ± SEM, n=143 cells for MGH-134 and n=174 cells for CALU6 were analyzed from 2 biological replicates) (see also Figure 3A). (D) Quantification of NADH/NAD^+^ ratio following NRF2 activation in KEAP1-independent cells. Cells stably expressing SONAR were treated and analyzed as described in (C) (Data are represented as a mean ± SEM, n=133 cells for H1975 and n=86 cells for HARA were analyzed from 2 biological replicates) (see also Figure 3A). (E-G) Depletion of KEAP1 in CALU6 (E) or MGH-134 (F) increases NADH/NAD^+^ ratio. NSCLC cell lines stably expressing SONAR and a DOX-inducible shRNA targeting KEAP1 or a non-targeting control were treated with DOX (100 nM) for 3 days and the NADH/NAD^+^ ratio was quantified as in (Data are represented as a mean ± SEM, n=171 cells for CALU6 and n=108 cells for MGH-134 from 2 biological replicates) (B). (H-I) Depletion of KEAP1 in H1975 or HARA (H) only moderately increases the NADH/NAD^+^ ratio. NSCLC cells stably co-expressing SONAR or indicated DOX-inducible shRNAs were treated and analyzed for changes in NADH/NAD^+^ ratio (I) as described in (E-F) (Data are represented as a mean ± SEM, n=179 cells for H1975 and n=131 cells for HARA were analyzed from 2 biological replicates). (J) NRF2 regulates the NADH/NAD^+^ ratio in KEAP1-dependent cells. Quantification of NADH/NAD^+^ in NSCLC cell lines expressing the indicated sgRNAs and treated with KI696 (1 µM) for 2 days (Data are represented as a mean ± SEM, n=241 cells for CALU6 and n=120 cells for MGH-134 were analyzed from 2-3 biological replicates) (see also Figure 3B). * indicates *p*-values < 0.05, ** indicates *p*-values < 0.01, *** indicates *p*-values < 0.0001. Statistical significance was determined by Stident’s t-test (B-D) and one-way ANOVA with Sidak’s correction for multiple hypotheses for (G-J). Scale bar: 25 µm.

**Figure S6:**
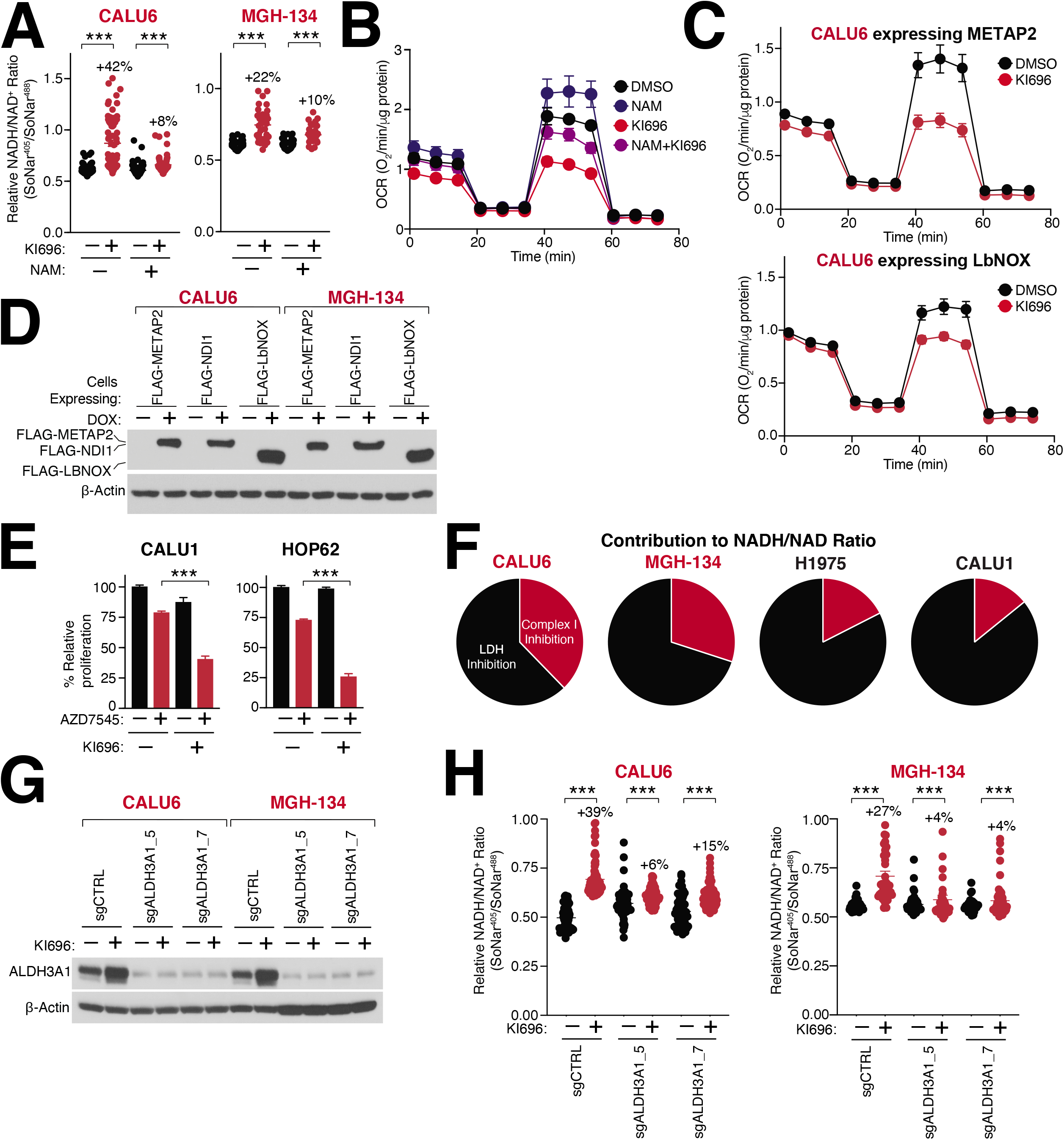
ALDH3A1 is a key regulator of the NADH/NAD+ ratio following NRF2 activation. (A) NAM treatment decreases the NADH/NAD^+^ ratio following NRF2 activation. NSCLC cell lines expressing SoNar were co-treated with NAM (1 mM) and KI696 (1 µM) for 2 days and the NADH/NAD^+^ ratio was determined by immunofluorescence analysis comparing the emission intensity after excitation at 408nm to 488nm (Data are represented as a mean ± SEM, n=210 cells for CALU6 and n=140 cells for MGH-134 were analyzed from 2 biological replicates) (**see also** Figure 3C). (B) NAM treatment rescues NRF2-mediated blockage of OCR. CALU6 cells were co-treated with NAM (1 mM), KI696 (1 µM) for 2 days or vehicle controls and OCR was determined by Seahorse Bioflux Analyzer (Data are represented as a mean ± SEM, n=6-8 biological replicates). (C) Over-expression of LBNOX overcomes NRF2-mediated blockage in OCR. CALU6 cells stably expressing DOX-inducible LbNOX or METAP2 (control) were pre-treated with DOX (100 nM) for 2 days followed by KI696 (1 µM) for another 2 days and OCR was determined as described in (B) (Data are represented as a mean ± SEM, n=6-8 biological replicates). (D) Representative immunoblot analysis of CALU6 and MGH-134 cells stably expressing DOX-inducible LbNOX, NDI1 or METAP2 following DOX treatment for 2 days. (E) Inhibition of pyruvate dehydrogenase kinase (PDK) sensitizes KEAP1-independent cells to NRF2 activation. NSCLC cell lines were co-treated with AZD7545 (PDK inhibitor, 100 µM) and KI696 (1 µM) for 6 days. Cell proliferation was determined by crystal violet staining (Data are represented as a mean ± SEM, n= 5 biological replicates). (F) KEAP1-depedendent cells have a greater reliance on Complex I for NADH oxidation. NSCLC cell lines expressing SONAR were treated with rotenone (Complex I inhibitor, 0.5 µM) followed by treatment with oxamate (LDHA inhibitor, 5 mM) and the change in NADH/NAD^+^ ratio was determined by flow cytometry comparing the emission intensity of 408 to 488. NADH/NAD^+^ ratio was normalized to the max ratio observed following treatment with both inhibitors (Data are represented as a mean ± SEM, n= 2 samples per group representative of two biological replicates). (G) Representative immunoblot of ALDH3A1 in CALU6 and MGH-134 cells expressing sgRNAs targeting ALDH3A1. (H) Depletion of ALDH3A1 in KEAP1-dependent cells reverts NRF2 mediated increase in NADH/NAD^+^ levels. NSCLC cell lines co-expressing SONAR and the indicated sgRNAs were treated with and KI696 for 2 days and the NADH/NAD^+^ ratio was determined by immunofluorescence analysis comparing the emission intensity after excitation with 408nm and 488nm (Data are represented as a mean ± SEM, n=429 cells for CALU6 and n=286 cells for MGH-134 were analyzed from 4 biological replicates) (see also Figure 3H). * indicates *p*-values < 0.05, ** indicates *p*-values < 0.01, *** indicates *p*-values < 0.0001. Statistical significance was determined by one-way ANOVA with Sidak’s post-hoc correction.

**Figure S7:**
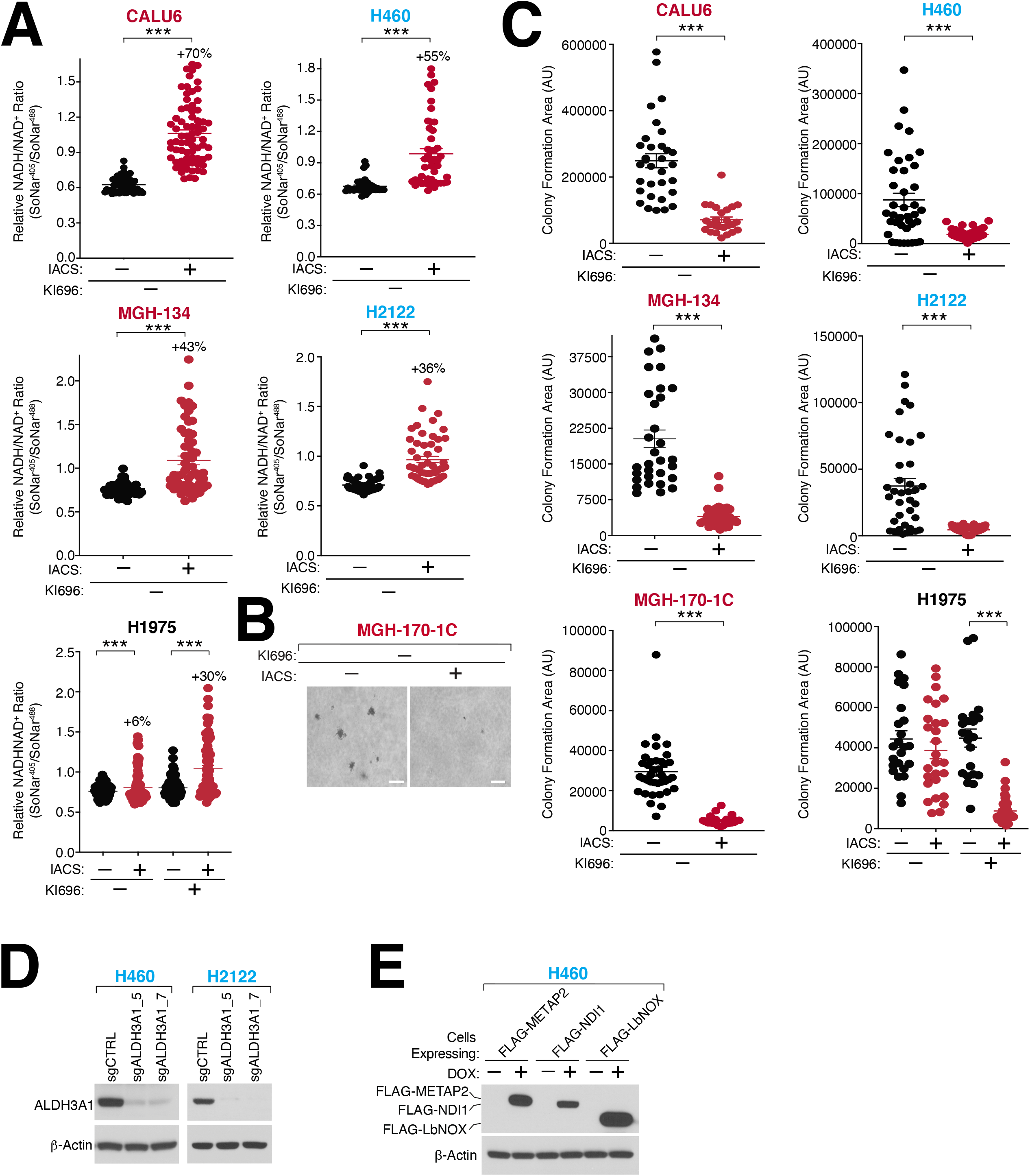
Increasing the NADH/NAD^+^ ratio selectively blocks the growth of NRF2 activated cells. (A) IACS-010759 (IACS) increases the NADH/NAD^+^ ratio in KEAP1-dependent and KEAP1-mutant but not KEAP1-independent cells. NSCLC cell lines expressing SONAR were co-treated with IACS (100nM) and KI696 (1μM) and the NADH/NAD^+^ ratio was determined by immunofluorescence analysis comparing the emission intensity following excitation at 408nm to 488nm (Data are represented as a mean ± SEM, n=129 cells for CALU6, n=100 cells for MGH-134, n=363 cells for H1975, n=188 cells for H460 and n=105 cells for H2122) (see also Figure 4A). (B) Representative images of MGH-170-1C cells grown in soft agar following treatment with IACS-017509 (200 nM). (C) Quantification of IACS treatment on NSCLC anchorage-independent growth. NSCLCs were co-treated with IACS (200 nM) and KI696 (1 µM) and soft-agar proliferation was determined by assessing colony forming area in multiple frames using ImageJ (Data are represented as a mean ± SEM, n= 4-6 biological replicates) (see also Figure 7C). (D) Immunoblot analysis of ALDH3A1 in H460 and H2122 cells expressing sgRNAs targeting ALDH3A1. (E) H460 cells stably expressing DOX-inducible FLAG-LBNOX, FLAG-NDI1 or FLAG-METAP2 were treated with DOX (100 nM) for 2 days and the expression of the indicated proteins was determined by immunoblot. * indicates *p*-values < 0.05, ** indicates *p*-values < 0.01, *** indicates *p*-values < 0.0001. One-way ANOVA and Student’s t-test were used to determine statistical significance. Scale Bar: 50 µm.

## Notes

### Competing Interest Statement

L.BP is a founder, consultant and holds privately held equity in Scorpion Therapeutics. D.E.F. has a financial interest in Soltego, a company developing salt inducible kinase inhibitors for topical skin-darkening treatments that might be used for a broad set of human applications. The interests of L.B-P and D.E.F. were reviewed and are managed by Massachusetts General Hospital and Partners HealthCare in accordance with their conflict-of-interest policies.

